# Single-cell transcriptomics reveals life cycle and cell cycle gene expression dynamics during *in vitro* growth of two *Leishmania* species

**DOI:** 10.64898/2026.09.29.755301

**Authors:** Catarina A. Marques, Ross F. Laidlaw, Felix Warren, Gabriel L.A. da Silva, Matthew Young, Kathryn Crouch, Richard Burchmore, Thomas D. Otto, Martin Llewellyn, Richard McCulloch, Emma M. Briggs

## Abstract

*Leishmania* species are transmitted between mammals by sand flies, within which they undergo a complex developmental programme involving transitions between replicative and non-replicative life cycle forms. Recent work using single-cell transcriptomics has revealed the developmental pathways of *Leishmania major* in the sand fly, but it is unclear how faithfully such development is recapitulated during *Leishmania* culture *in vitro*. In addition, no work has revealed how gene expression changes across the cell cycle of replicative *Leishmania* forms.

Using single-cell transcriptomics on unsynchronised cultures of *L. major* and *L. mexicana*, we addressed both questions. We show that *in vitro L. mexicana* promastigotes display the same two developmental endpoints seen in *L. major* cells recovered from sand flies, metacyclics and haptomonads. In contrast, in vitro *L. major* promastigotes development does not predict the presence of leptomonads or haptomonads, although non-dividing late metacyclics form through an early metacyclic stage as parasites enter stationary phase. Using computational reconstruction of the cell cycle in procyclic promastigote forms of both *Leishmania* species identified hundreds of genes whose transcript levels vary as the cell cycle progresses, revealing core conserved processes between the two species and with the closely related *Trypanosoma brucei*. These genes allowed us to examine connections between life cycle and cell cycle progression, revealing that some developmental transitions are strongly linked to a cell cycle phase, whereas others occur asynchronously. This work provides a comprehensive atlas and marker gene list for the interlinked cell and life cycles of two laboratory-cultured *Leishmania* species.

**Importance:** *Leishmania* parasites cause a group of neglected tropical diseases that can be fatal or cause life-altering lesions and are transmitted by biting sand flies. Much of our understanding of the complex developmental changes these parasites undergo within sand flies comes from laboratory-grown parasites, yet it is unclear whether these cultures accurately reproduce development. By examining gene activity in thousands of individual parasites, we discovered that two *Leishmania* species follow different developmental pathways under laboratory conditions, with only *L. mexicana* accurately recapitulating development in the sand fly. We also identified hundreds of genes whose activity changes during development and cell division. Some developmental transitions are closely linked to cell division, whereas others occur independently. Our results provide a detailed map of parasite development and markers for identifying different parasite forms. This resource will support future studies of how *Leishmania* parasites proliferate, develop, and prepare for transmission.

## Introduction

Over twenty species of the protozoan parasite *Leishmania* infect humans, causing between 700,000 and 1 million new leishmaniasis cases each year^1^. Forms of leishmaniasis vary in morbidity and mortality and are broadly grouped into cutaneous (CL) and mucocutaneous leishmaniasis (ML), the symptoms of which range from skin lesions/rashes to severe damage of the mucosal linings, and visceral leishmaniasis (VL), the fatal form of the disease. Although disease pathogenesis is directed by both the *Leishmania* species and host factors, all *Leishmania* species are spread between hosts by female sand flies (*Lutzomyia* or *Phlebotomus* genera) during a blood meal^2^. The parasite navigates varied extracellular and intracellular environments in the sand fly and mammal during this digenetic life cycle, as reflected in cellular plasticity and differentiation between life cycle forms, which have been mainly to date classified by their morphology^3,4^.

Metacyclic promastigote forms are injected by the sand fly into the mammal, where they are initially phagocytosed by neutrophils and macrophages and differentiate into non-motile amastigote forms^5^. Other host cell types, including fibroblasts, dendritic cells, epithelial cells and stem cells^6^, can also harbour amastigotes during infection, but macrophages appear to be the major location where amastigotes proliferate to increase parasitaemia. When ingested by a sand fly, amastigotes are released from host cells, allowing the change in temperature and pH to trigger differentiation into procyclic promastigote forms^7^. Various morphotypes of promastigotes then appear sequentially during growth in the sand fly, with the early ovoid procyclic forms being followed by elongated free-swimming nectomonad forms. These forms can cross the peritrophic matrix in the fly before migrating to the thoracic midgut. Here, long leptomonads develop before the appearance of haptomonad and metacyclic forms completes the life cycle^8,9^. Notably, early morphology-based studies demonstrated that the appearance and the length of persistence of each form varies between species, yet the sequential order appears to be conserved at least between *L. mexicana* and *L. infantum* ^4^.

Initial quantification of post mitosis (post-M) parasites that have duplicated and divided their nuclear and mitochondrial (kinetoplast) genomes, but have not yet undergone cytokinesis, demonstrated metacyclic forms to be non-proliferative *in vitro* and *in vivo*, as no post-M cells could be identified for this population. Additionally, only low proportions of post-M phase nectomonad forms were observed (0.1-4.9 %), indicating nectomonads are also a largely non-replicative stage, in contrast to procyclic (6-65.6 % post-M) and leptomonad (4.7-13.5 % post-M) forms^4^.

Interestingly, percentages of post-M phase procyclics vary significantly between *L. infantum* (6.0-9.9%) and *L. mexicana* (42.4-56.5%) *in vitro* and *in vivo*. The varied morphologies of *L. mexicana* and *L. major* promastigotes cultured *in vitro* are also linked to cell cycle progression^10–12^. Cell body length increases throughout G1 before shortening post-S phase^11,12^. This wide range in cell lengths does not change with increasing cell culture density (3×10^6^ – 1.3×10^7^ cells/ml). In log phase promastigotes, cell division results in the generation of two daughter cells, one with a new, short flagellum and the other with a longer, old flagellum; as these lengthen during subsequent G1 phases, average flagella length across the population increases with cell density. Increasing cell width does not correlate with flagella length, but cell size and morphology are significantly altered post-S phase^11,12^. Given this mix of cell sizes and shapes, defining cell cycle phases relative to life cycle forms *in vitro* is complex based on morphology alone.

Very recently, developmental life cycle forms have been resolved for *L. major* in the sandfly using transcriptomic signatures captured with single cell RNA-seq^13^. This work demonstrated that, like *L. mexicana* and *L. infantum, L. major* also sequentially develops from procyclic to nectomonad, then to leptomonad and finally metacyclic forms^13^. Trajectory inference additionally placed haptomonads as an alternative developmental endpoint that, like metacyclics, develop from the leptomonad population. Notably, haptomonad forms, like metacyclic forms, were demonstrated to support and even establish infections in mice and make up the majority of the promastigote population egested by infected sandflies, highlighting their important role in infection. A likely replicative metacyclic precursor stage in *L. major* was also detected, which was not identified in early morphology-based studies of *L. mexicana* or *L. infantum*^4^.

In this study, we have used scRNA-seq to understand the diversity of promastigote forms present in *in vitro* cultures of *L. mexicana* and *L. major* species, both causative agents of CL in humans. Both species are commonly studied due to the ease of culture, availability of genetic manipulation tools and, in the case of *L. mexicana*, ability to recapitulate promastigote to amastigote differentiation *in vitro*^3^. What is unclear, however, is to what extent parasite development *in vitro* follows what is known in the sand fly. Moreover, cell cycle gene expression changes or how these might link to life cycle transitions have not yet been explored. Here, we find that cultured *L. mexicana* promastigotes effectively reproduce the branched development from procyclic forms to both metacyclic and haptomonads end points. However, we do not detect haptomonads in *L. major* cultures, where only metacyclics developed. Further analysis of these scRNA-seq datasets allowed us to uncover the cell cycle-regulated transcriptome of each species’ procyclic forms. Comparison with the cell cycle regulated genes of *Trypanosoma brucei* procyclic forms identified conserved transcriptional markers for G1 and S phase but highlighted greater variation in transcripts that define G2/M phase cells between species. Combining these cell cycle markers with the previously defined life cycle markers^13^ allowed us to understand the cell cycle and life cycle relationship of promastigotes, revealing both cell cycle phase-linked and asynchronously occurring developmental transitions.

## Results

### Identification of life cycle forms in unsynchronised *in vitro* cultured *L. major* promastigotes

*L. major* promastigotes, like other *Leishmania* species, display a range of morphologies reflecting the mitotic cell cycle when growing *in vitro*^10^, but the extent to which these morphologies may additionally reflect the different sand fly forms seen *in vivo*^4,13,14^ is unclear. To ask if single-cell transcriptomics can address these limitations, we captured populations of *L. major* that were exponentially growing (Log) or had just entered ‘stationary phase’ (STA) after 72 hrs of culture (Fig. 1A). Consistent with previous work^15^, flow cytometry showed that while Log cells had the range of DNA content expected for cycling cells, STA cells had substantially reduced 4N (G2-M phase) and 2N-4N (S phase) DNA content and a concomitant increase in 2N (G1 phase), indicating enrichment for non-replicating cells (Fig.1B). These populations were then each multiplexed with *T. brucei* cells^16^ and subjected to scRNA-seq using the Chromium single cell 3′ workflow (10X Genomics) and Illumina sequencing in a single batch. Transcriptomes were filtered to remove doublets and lower quality transcriptomes (using levels of rRNA and mitochondrial transcripts), as described previously^17^. After filtering, we recovered similar numbers of individuals of

**Figure 1.**
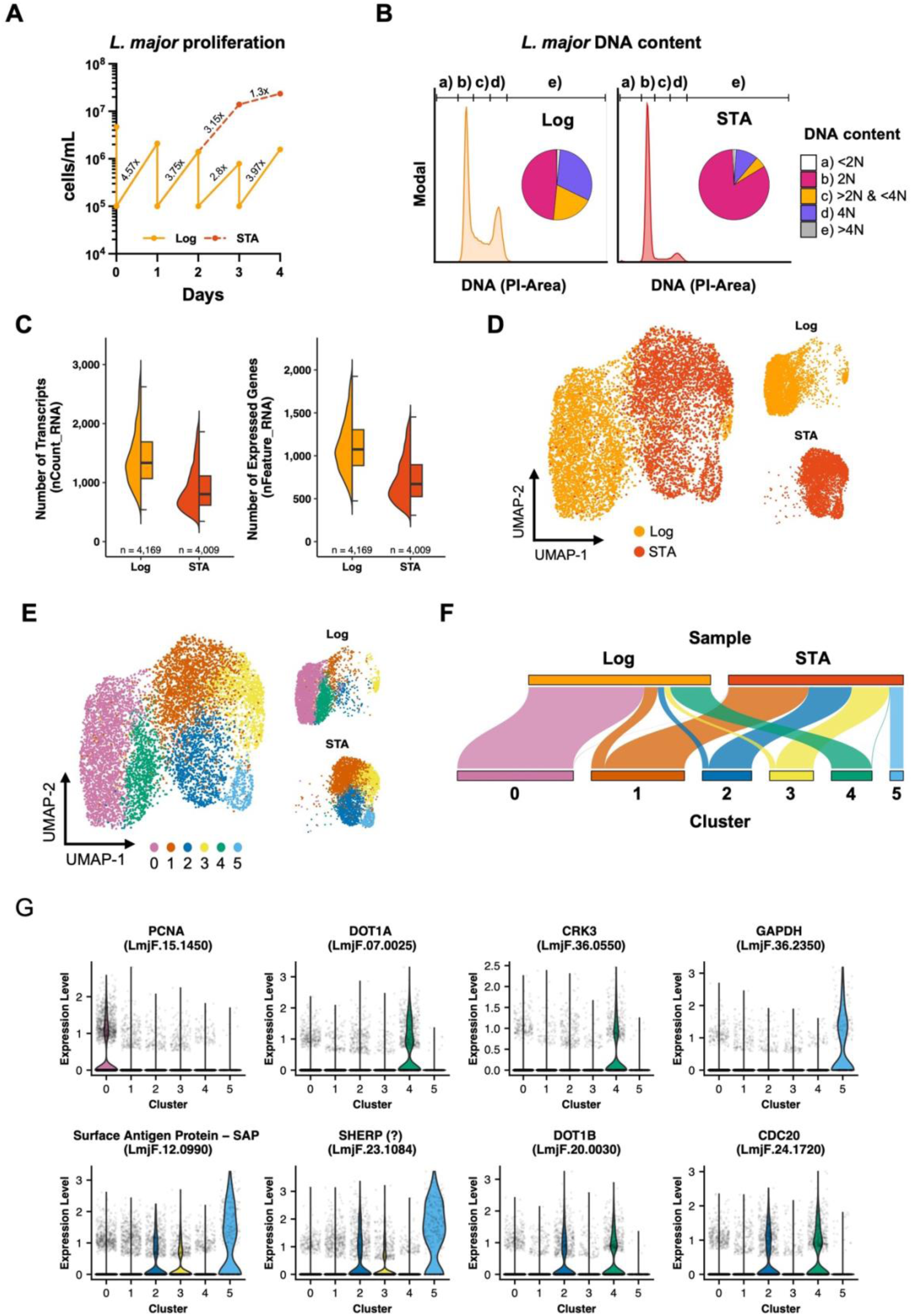
Single cell transcriptomic analysis of *Leishmania major* promastigotes in culture. **A)** Growth curve of *L. major* promastigote forms. Log phase sample was diluted to 1×10^5^ cells/ml every 24 h, whereas the stationary (STA) phase sample was allowed to reach maximal density after 48 h. Numbers indicate number of division cycles. **B)** Flow cytometry analysis detecting DNA content by propidium iodide (PI) staining. Insert pie charts show the percentage of cells with a DNA content of 2N (proxy for G1 phase), >2N <4N (S phase), 4N (G2M phase), as well as cells with <2N and >4N DNA content. **C)** Violin plots show the range of number of transcripts (left) or number of genes (right) per cell. Log sample represented as dark yellow; STA sample as dark orange. n indicates the number of cells in each dataset after QC filtering. **D)** UMAP of the merged Log and STA single cell transcriptomes, coloured by sample. Smaller UMAPs (right) show the sample split, for a better visualisation of the two samples. **E)** UMAP of the merged samples coloured by the identified clusters. Smaller UMAPs (right) show the sample split, for a better visualisation of the two samples. **F)** Alluvial plots showing the proportion of each sample assigned to each cluster. **G)** Violin plots of selected cluster markers examples.

*L. major* log (4,169) and stationary (4,009) transcriptomes; however, the median number of transcripts and genes detected per cell in the Log sample was notably higher (1,334 transcripts; 1,074 genes) than that of the STA sample (804 transcripts; 671 genes) (Fig. 1C, Fig. S1A). The mean transcripts per cell recovered from Log parasites (1,413) is comparable to previous analysis of *in vitro* grown *L. tropica*^18^ also in Log phase growth (1,079 mean transcripts per cell^18^). The lower number of transcripts per cell in our STA sample may reflect lower overall RNA content in STA cells (Fig.1C) relative to Log cells, as has been previously observed in *L.* (*Viannia*) *lainsoni* promastigote forms^19^.

Data from the Log and STA samples were normalised, merged, visualised as a UMAP (Uniform Manifold Approximation and Projection) via dimensionality reduction (Fig. 1D) and subjected to unsupervised cell clustering (Fig. 1E), resulting in six clusters across the two samples (Fig. 1E, F). Two clear clusters (clusters 0 and 4) made up the majority of the Log *L. major* sample (∼84.6%), with a minority of cells from this sample grouping in clusters 1, 2 and 3 (∼14.3%). In contrast, the STA parasite transcriptomes largely grouped to these three latter clusters (1, 2 and 3, ∼76.8% of the cells), as well as a smaller cluster 5 (∼23.2%), which is largely unique to the STA sample (306/309 cells ; Fig. 1E, F). To begin to determine the potential biological differences between the clusters, we first identified the most differentially expressed genes (i.e. cluster markers) in each cluster relative to the others (Fig.S1B Supplementary data 1). The most enriched transcript in cluster 0 was proliferating cell nuclear antigen (PCNA; Fig. 1G, Fig. S1B), whose expression is most prominent in G1/S phase of *L. donovani* cells, where it co-localises with signals of DNA replication in the nucleus^20^. In the other Log-specific cluster (4), a highly abundant transcript detected was DOT1A (Fig. 1G, Fig. S1B), whose activity has not been analysed in *Leishmania* but plays a crucial role in mono- and di-methylation of histone H3K76 during DNA replication in *T. brucei*^21^, where these histone modifications are limited to G2/M-C phase cells and are distinguishable from PCNA expression, which peaks in S-phase^21,22^. Furthermore, a second prominent transcript detected in cluster 4 was cdc2-related kinase 3 (CRK3) (Fig. 1G, Fig. S1B), whose conditional deletion in *L. mexicana* leads to G2/M phase arrest^23^. These data are consistent with clusters 0 and 4 being distinct populations of replicating cells (explored in more detail below). Amongst the four STA-enriched clusters, cluster 5 was marked by high level of expression of GAPDH and substantial expression of SAP and SHERP (Fig. 1G, Fig. S1B); expression of GAPDH in some *Leishmania* species has been associated with resistance to nitric oxide^24,25^, while SHERP is needed for metacyclogenesis^26^ and shows increased expression in both non-dividing nectomonads and metacyclics^27^. While these genes do not reveal a discrete identity for cluster 5, DOT1B showed notably increased expression (Fig. 1G, Fig. S1B) in clusters 2 (STA-rich) and 4 (Log-rich). In *T. brucei*, DOT1B is essential for differentiation of bloodstream form cells to procyclic^28,29^ and although loss of the methyltransferase does not impede transition from Log to STA phase of *L. mexicana* grown *in vitro*, or differentiation of *L. mexcicana* promastigotes to amastigotes *in vitro* or *in vivo*^30^, these data hint at a developmental progression from replicating to non-replicating *L. major* cells *in vitro*. Such progression may be linked to cell cycle progression, given the very similar elevated levels of CDC20^31,32^ expression in clusters 2 and 4 (Fig.1G, Fig. S1B). The highest expressed marker genes of cluster 3 were histones (Fig. S1B), and there was a lack of highly expressed genes unique to cluster 1, confounding attempts to further define these STA sub-populations by this approach alone (Fig.S1B).

Biological Process GO terms enrichment analysis revealed that clusters 0 and 4 share enrichment of multiple terms (Fig.S1C, Supplementary data 1) relating to glucose metabolism and ATP production, consistent with these clusters being actively proliferative forms. Cluster 1 markers were notably not related to particular GO terms, with recovered terms relating to only single genes (Supplementary data1), including transport due to the expression of Aquaglyceroporin 1 (AQ1), which locates to the flagellum in *L. major*, but to the glycosomes in other *Leishmania* species^33^.

Clusters 2 and 4 shared upregulation of genes relating to microtubule-based movement, indicative of cell division processes or cytoskeleton remodelling (Fig. S1B). Cluster 3 GO terms related to translation and are represented by ribosomal proteins and other translation factors, while cluster 5 GO terms may relate to a switch in metabolic process, as gluconeogenesis genes were upregulated. Together, these data hint at the identification of life cycle transitions occurring during *L. major in vitro* culture.

### The cell cycle regulated transcriptome of *L. major* promastigotes reveals that stationary phase populations contain a mix of proliferative and arrested forms

We next sought to determine the cell cycle regulated transcriptome in *L. major,* focusing first on the Log *L. major* promastigote sample, as this consists of proliferating forms (Fig 1A,B), providing a high confidence dataset to infer cell cycle dynamics. To this end, we first took advantage of our previously published scRNA-seq data in *T. brucei*^16^, which revealed transcripts with expression changes across the cell cycles of both bloodstream and procyclic form cells. From the *T. brucei* procyclic form gene list that includes clear markers of G1, S and G2/M phases, we identified 615 *L. major* orthologues. The gene list was further slimmed by removing gene duplicates and hypotheticals, leaving 213 cell cycle phase marker genes (Supplementary data 2). These markers were then used to label the cell cycle phases of both the Log and STA *L. major* samples (Fig. 2A), as described in Briggs et al 2023. As expected, this labelling reflected the nature of the two samples: ∼94.4% of the Log cells were labelled as G1, S or G2/M and only ∼5.6% were classified as “unlabelled”, meaning no phase marker set was highly expressed above background levels; in contrast, ∼52.4% of the STA cells were classed as “unlabelled” and ∼47.6% were labelled as being in one of the cycling phases (Fig. 2A; Fig. S2A).

**Figure 2.**
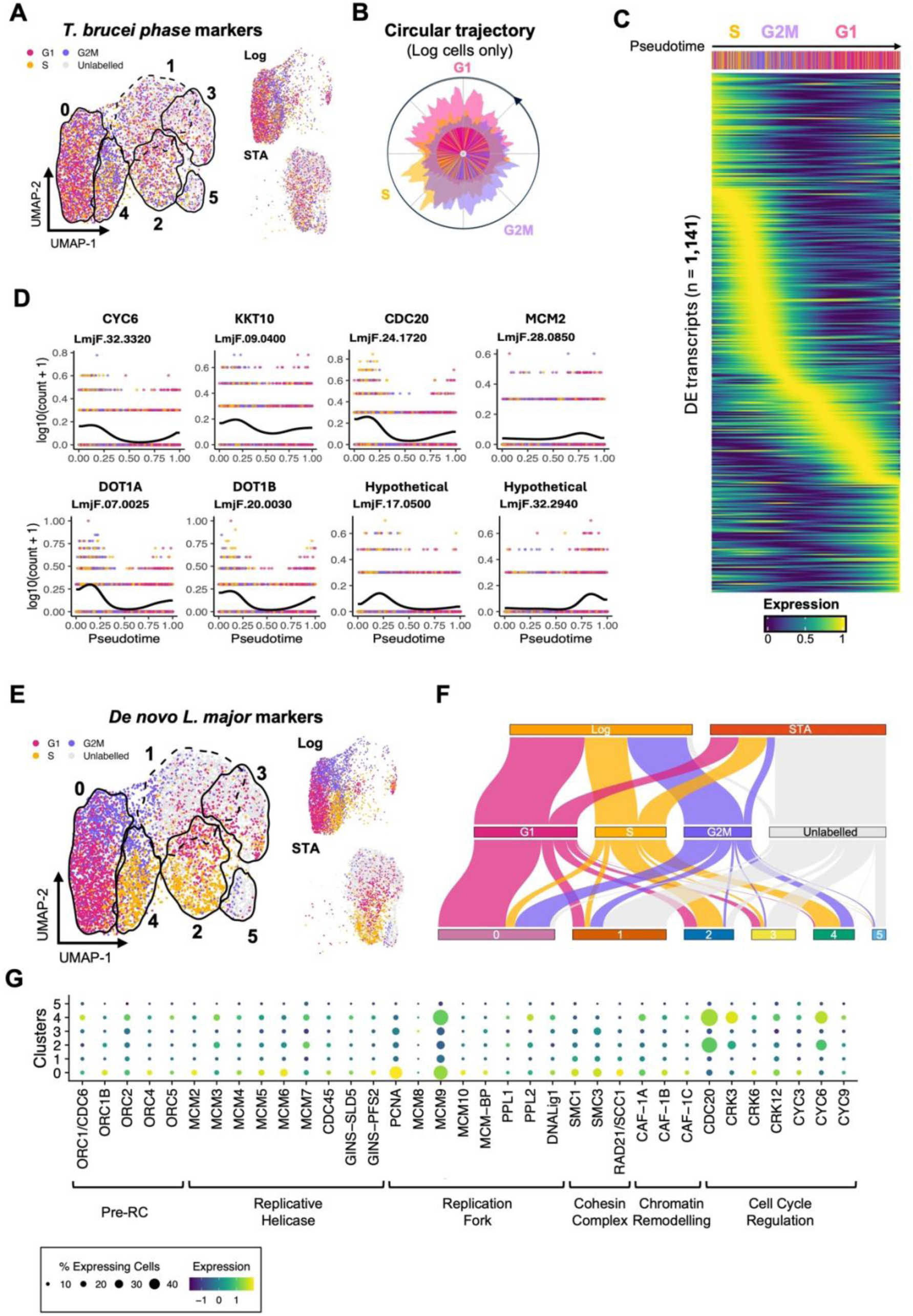
Defining the *L. major* cell cycle regulated transcriptome. A) UMAP of *L. major* promastigotes coloured by cell cycle phase using phase markers identified previously in *T. brucei*^16^. Smaller UMAPs (right) show the sample split, for a better visualisation of the two samples. **B)** Cyclum generate circular trajectory of Log sample promastigotes coloured by cell cycle phase. The centre shows individual cells coloured by phase (as in A), while the outer ring shows 3 histograms, one per phase, of the number of cells. **C)** Heatmap showing scaled expression (z-score) of significantly differentially expressed genes over the cell cycle. **D)** Log10 transcript counts of selected genes over pseudotime. Cells are coloured by phase, while the black line represents the average expression pattern across pseudotime. **E)** Both Log and STA samples coloured by cell cycle phase using *L. major* markers filtered from the genes in C). Each cluster is delineated by a black line on top of the UMAP and numbered accordingly. Smaller UMAPs (right) show the sample split, for a better visualisation of the two samples. **F)** Alluvial plot showing the proportion of each sample (top) and cluster (bottom) labelled as each cell cycle phase (middle). **G)** Dot plot showing expression of known cell cycle factors across each cluster.

Overall, these results roughly match the flow cytometry data (Fig. 1B) but suggest that the cell population in the STA sample containing 2N DNA content includes both G1 and apparently non-cycling cells (unlabelled, Fig. S2A). Nevertheless, the cells did not clearly group together by cell cycle phase (Fig. 2A).

To identify *L. major*-specific cell cycle regulated genes and, consequently, obtain *Leishmania-*specific markers for cell cycle phase labelling, we took a periodic trajectory inference approach (Cyclum^34^) using only the Log sample cells (Fig. 2A, top right). This approach assigns pseudotime values to the cells, independently of the previously allocated cell cycle phase labelling. The resulting circular trajectory was then plotted using the previously allocated cell cycle phase labels to assess success in capturing the cell cycle (Fig. 2B). PseudotimeDE^35^ was then used to identify differentially expressed genes across pseudotime, reflecting the cell cycle (Fig. 2C). Using a false discovery rate (FDR) adjusted p-value of < 0.05 and mean log (fold change) of > 1.5, a total of 1,141 genes with significant cell cycle-associated changes in transcript abundance were identified (Fig. 2C), along with enriched GO terms associated with these genes (Supplementary data 2). As expected, GO terms included terms linked to the flagellum and cell motility, translation, cell cycle and organelle assembly. Genes with clear roles in the cell cycle included cyclin 6 (CYC6, LmjF.32.3320), cdc2-related kinases CRK2 (LmjF.05.0550) and CRK3 (LmjF.36.0550), CDC20 (LmjF.24.1720), and kinetochore protein KKT10 (LmjF.09.0400). Moreover, five components of the DNA replication helicase (MCMs 2, 3, 5, 6 and 7) were also differentially expressed, as were DOT1A and DOT1B, and 469 genes encoding hypothetical proteins that were expressed at various points in the cell cycle (Fig. 2D, Supplementary data 2).

By applying a mean log (FC)>2 threshold to the above list of 1,141 cell cycle-regulated genes, we robustly defined new *L major*-specific markers for G1 (147 genes), S (202 genes) and G2/M (127 genes). These “*de novo*” *L. major* markers were then used to re-assign a cell cycle phase to each individual cell in both the Log and STA samples (Fig. 2E). Comparing these assignments to the previous labelling using the *T. brucei* cell cycle markers (Fig. 2A), there was now clear groupings of cells in G1, S and G2/M (Fig. 2E), and the proportions of cells in the Log sample labelled as G1 (40.9%), S (23.4%) and G2/M (32.6%) (Fig. 2F; Fig. S2B) compare closely with the flow cytometry data (Fig. 1B; ∼48.3% 2N cells, ∼19.4% >2N and <4N cells, and ∼30.7% 4N cells). Regarding the STA sample, most of the cells remained classed as unlabelled (from ∼52.4% to 63.2%), while the proportion of cells classed as S increased slightly (from 12.7% to 16.1%), and the number of cells labelled as G2/M decreased from 17.7% to 4.6% (Fig. S2A, B). This again suggests that the 2N DNA content cells detected by flow cytometry include both G1 and non-cycling cells (Fig. 1B), while the number of cells with an “S phase” or “G2/M phase” transcriptome is not entirely reflected by, respectively, >2N and <4N or 4N DNA content cells (∼5.1% and 10.23%, respectively; Fig. 1B).

A more robust labelling of the cells now allowed for a deeper analysis of the previously identified (Fig. 2E, F). Most cells in clusters 0 and 4 (∼95.7% and ∼98.7%, respectively), which originate almost entirely (>98%) from the Log sample, were predicted to be in either G1, S or G2/M phases of the cell cycle. However, the phase distributions were different between the two clusters: ∼57.8% of cluster 0 was labelled as in G1 phase, ∼8% of the cells were S, and ∼29.9% were G2/M phase; in contrast, in cluster 4 only ∼4.2% of the cells were labelled as G1, ∼61.7% of the cells were S, and ∼32.8% G2/M (Fig. 2F; Fig. S2B). Together, these observations are consistent with the cell cycle associated markers that were identified for these clusters above (Fig. 1G, Fig. S1C). In the case of clusters 1, 3 and 5 (where most originate from the STA sample; Fig.1F), the proportion of unlabelled cells was, respectively, ∼67.5%, ∼66.2% and 79.3% (Fig. S2). In all three clusters the percentage of cells in S phase ranged from 4.5-9% (Fig. S2B), while the percentages of cells in G1 and G2/M phases varied from 2.9-20.3% (Fig. S2B). In the case of cluster 2, ∼53.2% of the cells were labelled as S, ∼25.8% as G1, 5% as G2/M and ∼16% as unlabelled (Fig. 2F; Fig. S2B), indicating that this is a cluster composed of mostly cycling cells. Overall, these data are consistent with previous work that indicated DNA replication is generally suppressed when *L. major* enters stationary phase^15^, although the scRNAseq now reveals that some STA cells do remain cycling. Indeed, analysis of the expression of a wide range of *L. major* DNA replication and cell cycle regulation-associated factors reveals that expression of nearly all of these are downregulated in STA clusters 1, 3 and 5 relative to Log clusters 0 and 4 (Fig. 2G).

### *Leishmania major* grown in culture does not form all life cycle stages seen in the sand fly

To understand the potential links between the cell and life cycles of *L. major* (Fig. 3A), we next analysed our *in vitro* samples using life cycle form-specific markers previously identified from scRNA-seq of *L. major* recovered from *Phlebotomus duboscqi*^13^. For each life cycle stage, we first selected 5 genes from the life cycle markers defined *in vivo* that were also detected in our samples and analysed their expression across the six detectable clusters (Fig. 3B). Next, to better understand the nature of our *in vitro* samples’ clusters, we expanded the analysis to a larger number of *in vivo* markers and mapped their average expression onto our *in vitro* samples UMAP (Fig. 3C, D). This analysis revealed that clusters 0 and 4 are, as expected, proliferative procyclic forms, with cluster 0 having high expression of procyclic, G1 and G2/M markers genes (Fig. 3D), and cluster 4 high expression of procyclic, S and G2/M markers (Fig. 3D). Nectomonads were discernible as cluster 3, which is found in the STA-rich population, and consistent with pronounced expression of the protein kinase NEK15 in these forms (Fig. S1B). Nectomonads are believed to be non-proliferative^2,24^, which is supported by our analysis, as cluster 3 did not show expression of G1, S or G2/M markers (Fig. 3D). In our *in vitro* data a clearly distinct transitional cluster between procyclics and nectomonads was not obvious (Fig. 3C, procyclic to nectomonad), but markers genes relating to this transition were expressed in cells spatially close to labelled nectomonad cells.

**Figure 3.**
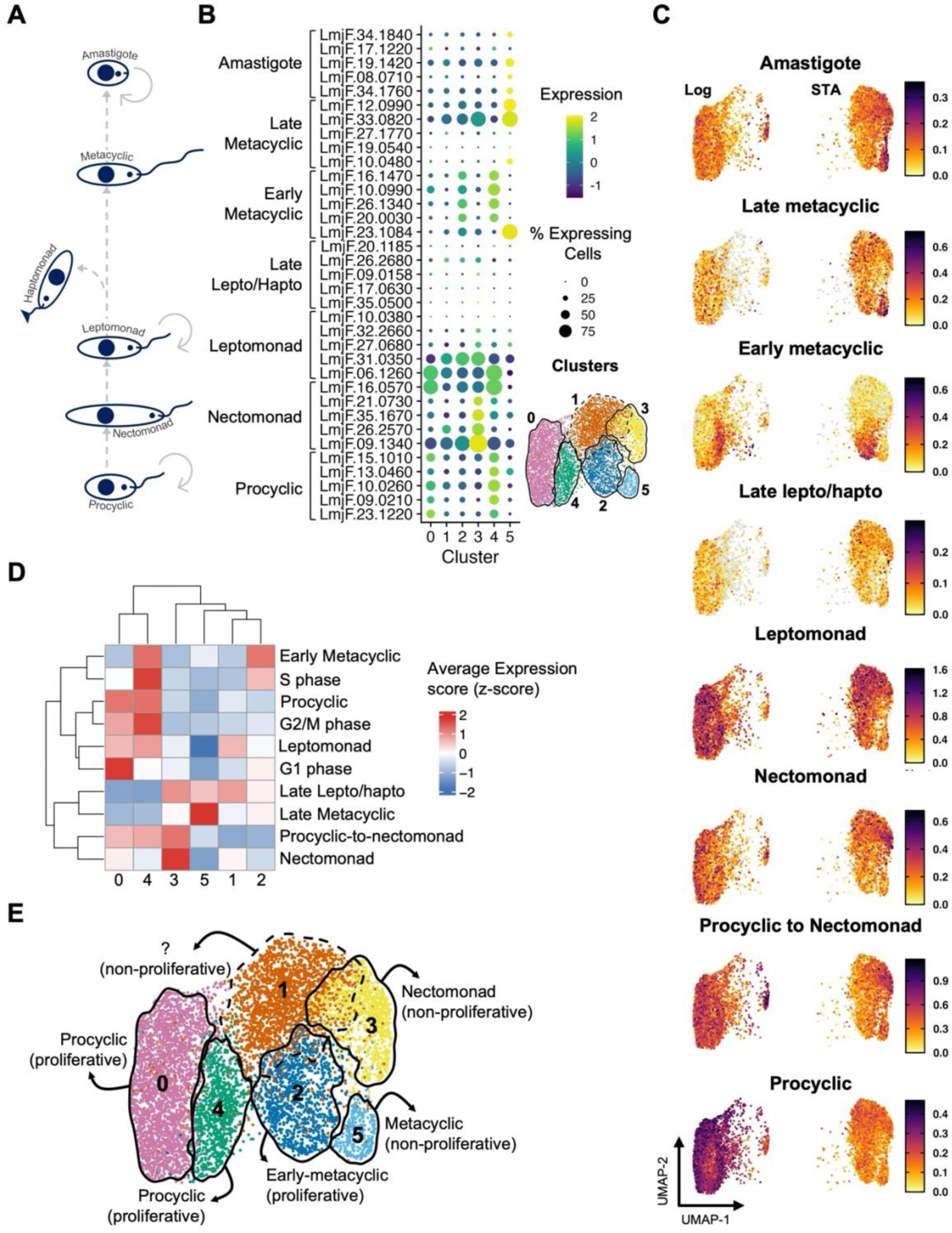
Defining the life cycle forms of *Leishmania major* promastigotes in culture. **A)** Diagram showing the prevailing model of *Leishmania* life cycle stages prior to scRNA-seq studies, based on Cecílio *et al.* (2022)^2^ and Dostálová and Volf (2012)^88^. Not to scale. **B)** Dot plot indicating expression of marker genes identify by Catta-Preta et al. for *L. major* clusters (shown to the right for reference). **C)** UMAPs of log and STA samples coloured by average expression scores of complete marker list identified previously for each life cycle form. Scores were calculated as average marker expression per score, relative to the average expression of a random control set of genes. **D)** Scaled expression scores for both life cycle scores shown in C) and cell cycle phase scores, for each cluster with clustering to identify relationships. **E)** UMAP of *L. major* coloured by cluster, with cell type annotations.

Nectomonads give rise to replicating leptomonads (Fig. 3A), which *in vivo* scRNA-seq revealed to be the progenitors for either non-replicating haptomonads or metacyclics^13^. Here, there was no convincing gene expression evidence for the presence of either late leptomonads or haptomonads in our *in vitro L. major* samples (Fig. 3B, C). Although leptomonad markers were expressed, this was without clear enrichment to a specific cluster (Fig. 3B, C), instead displaying expression at higher levels in clusters 0 and 4 (procyclic) and cluster 1 (Fig. 3D). These data may suggest cluster 1 comprises leptomonad-like forms, but there was no expression of cell cycle phase markers in this cluster (Fig. 3D, Fig. 2E, F); furthermore, the equal enrichment of leptomonad markers in cluster 1 and in the two procyclic form clusters (0 and 4) was not seen in *L. mexicana* (see below). Neither individual marker genes (Fig. 3B) nor combined expression of the complete list of markers (Fig. 3C) provided evidence for late leptomonad/haptomonad in any cluster, indicating that these do not form, at least in the *in vitro* culturing conditions used here. Despite this absence, cluster 5 showed elevated expression of markers of late metacyclic forms, while their presumed progenitors (‘early metacyclics’) were mostly consistent with cluster 2, a cluster with evidence for being replicative (Fig. 3D), as they share expression of several markers also seen in cluster 4, including DOT1B and DNA ligase k alpha (Fig. S1B) and clearly display G1 and S transcript profiles. Consistent with our experimental approach, where no amastigotes should be present, amastigote markers were most expressed in late metacyclics (cluster 5), reflecting pre-adaptation to survival in the mammal.

Taken together, the above scRNA-seq data reveal several aspects of the growth of *L. major* promastigotes in culture. Exponentially growing *L. major* populations are dominated by replicating procyclics and these are highly distinct from early stationary phase cells that have remained above 1×10^7^ cells/ml for just 24 h. This STA population is surprisingly heterogenous and contains at least two discernible non-replicating forms (nectomonads and late metacyclics). Amongst the two other populations seen in stationary phase, one appears to be a progenitor of metacyclics (cluster 2), as seen *in vivo*^36^, while the other (cluster 1; Fig.1B) could not be confidently identified as, despite expressing leptomonad-associated transcripts, there was no evidence to support replication of this cell type. Interestingly, amongst the most prominently expressed gene in cluster 1 was HAP2 (Fig. 1F), which encodes a conserved meiosis factor that has been implicated in *Leishmania* mating^18,37^. Despite this range of putative life cycle forms, *L. major* grown *in vitro* (at least in the conditions used in this study) does not appear to fully recapitulate the forms seen *in vivo* (Fig. 3E), since in addition to the apparent absence of replicating leptomonads we could not clearly detect non-replicating haptomonads, suggesting the absence of this developmental branch. Though this may simply be due to inefficient capture of these life cycle stages in our *L. major* samples, it is notable that these forms were readily detectable in *in vitro* growing *L. mexicana* populations (see below).

### The developmental trajectory of cultured *L. mexicana* promastigotes closely resembles that of *L. major* in the sand fly

In contrast to *L. major*, which has been reported to be refractory to efficient induction of differentiation to axenic amastigotes *in vitro*^38^, *L. mexicana* efficiently undergoes such differentiation^39^. For this reason, we expanded upon the approach we took in *L. major* by growing *L. mexicana* promastigotes from 1×10^5^ cells.mL^-1^ for 7 days without passaging, sampling approximately 3,000 cells at each of 24 h, 48 h and 144 h (Fig. S3A). Axenic amastigotes were obtained by transferring stationary phase promastigotes cells from HOMEM media into cSDM media (pH 5.5), and incubating them at 32 °C with 5% CO_2_, starting at a concentration of 1×10^6^ cells.mL^-1^. Samples of 3,000 cells were then taken at 24 h and 72 h post-transfer (Fig.S3B). The experiment was performed in biological duplicates: in the first replicate all five *L. mexicana* timepoints were combined in equal numbers into a single sample for scRNA-seq (sample 1); for the second independent replicate, we split these time points into two scRNA-seq samples containing either promastigote (24 h, 48 h, 144 h; sample 2Pro) or axenic amastigote (24 h and 72 h; sample 2Ama) timepoints (Fig. S3C). In each case, samples were multiplexed with *T. brucei* or *T. congolense* ^16,17^, and subjected to scRNA-seq as described above. After quality control and filtering as described above for *L. major*, the three *L. mexicana* samples were combined by first merging samples 2Pro (promastigote only) and sample 2Ama (amastigote only), and then by integrating the merged data with sample 1 (promastigotes and amastigotes) using Seurat’s CCA integration method to remove potential batch effects between the replicates (Fig. S3D, E). This combined dataset was then subjected to unsupervised cell clustering and visualised as a UMAP (Fig. S3F, Supplementary data 3). As with *L. major*, the expression levels of previously identified life cycle makers were also plotted to identify likely cell types in these data (Fig. S3G).

Initial clustering analysis identified 8 clusters (Fig. S3F). This included cluster 1, which contained parasites only found in the amastigote culture conditions (samples 1 and 2Ama), with virtually no cells from promastigote conditions (sample 2Pro).

Despite this, the transcriptomic signature of cluster 1 was most like that of procyclic and leptomonad forms, not amastigote forms (Fig. S3G). In our experimental set up a mix of promastigote forms were transferred to amastigote differentiation conditions; therefore, this cluster is most likely promastigote forms that have persisted in the short exposure to these conditions but are unlikely to represent a biologically meaningful cell type. We therefore removed this cluster from further analysis (Fig. 4A).

**Figure 4.**
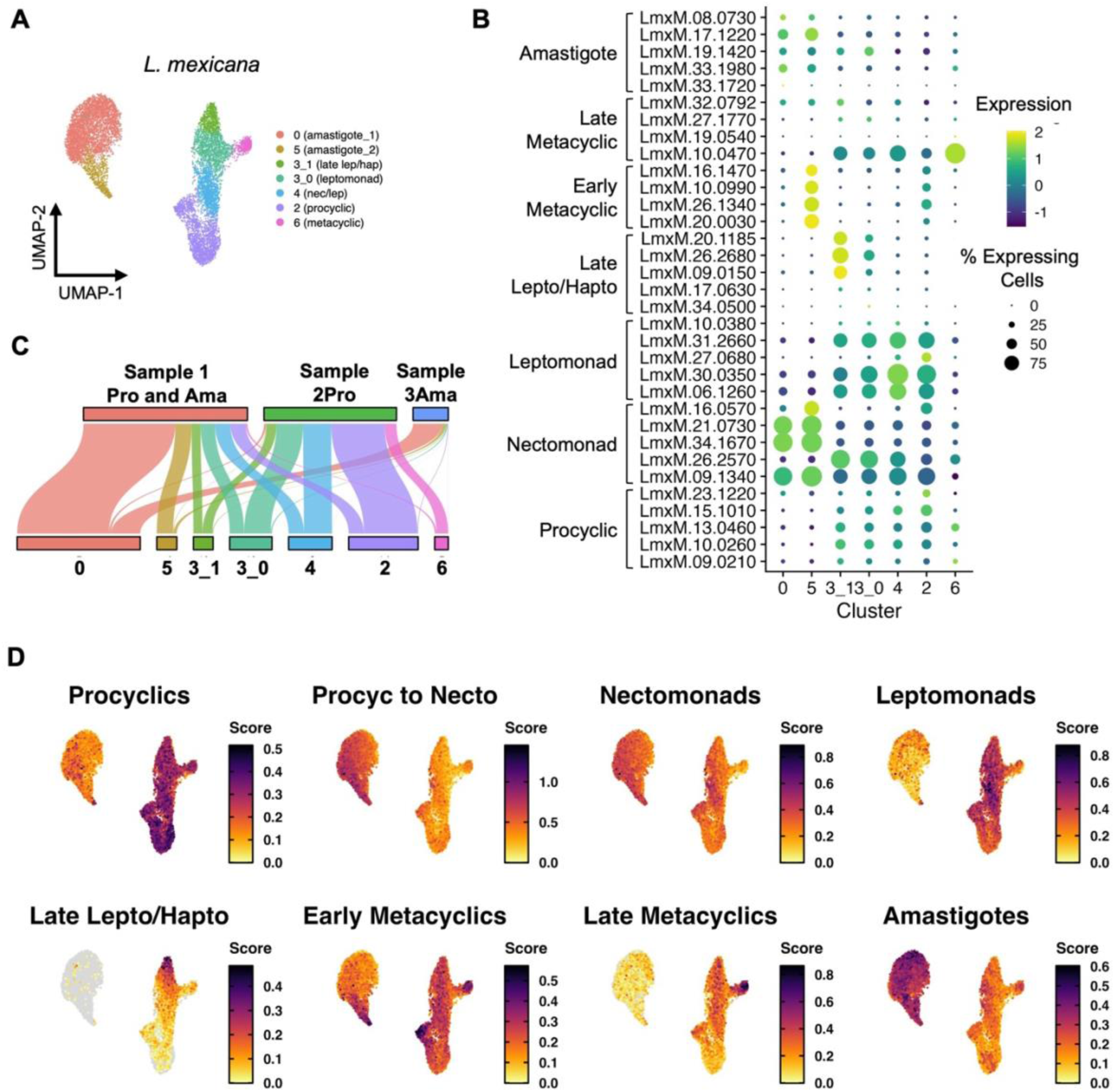
Life cycle progression of *Leishmania mexicana* from promastigotes to amastigotes in culture. **A)** UMAP of *L. mexicana* coloured by cluster and life cycle form. **B)** Dot plot indicating expression of homologous marker genes identify by Catta-Preta et al. for *L. major* clusters. **C)** Proportion of each sample assigned to each cluster shown in A). **D)** UMAPs of the *L. mexicana* samples coloured by average expression scores of complete homologous marker list identified previously, for each life cycle form.

The remaining clusters (Fig. 4A, C) were revealed as corresponding to most of the *L. major* life cycle forms described in sand flies^13^, both by examining average expression of life cycle marker gene sets to the UMAP (Fig. 4D) or analysing expression of the 4-5 life cycle form-specific markers (Fig.4C). These analyses clearly indicated a late leptomonad/haptomonad population (missing in *L. major*) within cluster 3 (Fig. 4B, D). We therefore subclustered this group into 3_0 and 3_1 to distinguish leptomonads and late leptomonad/haptomonad, as described by Catta-Pretta et al^13^. This resulted in seven cell types with distinct transcriptomic signatures (Fig. 4B, C), including clear amastigote (clusters 0 and 5), procyclic (cluster 2), late leptomonad/haptomonad (cluster 3_1) and late metacyclic (cluster 6) clusters. As with *in vitro L. major*, an *L. mexicana* cell population transitioning between procyclics and nectomonads^13^ was not apparent *in vitro*. *L. mexicana* nectomonad and leptomonad populations were not distinct from one another based on transcriptomic profile, but transcript markers of both forms were clearly expressed and mostly localised to clusters 4 and 3_0 (Fig. 4B, D). Notably, in contrast to *L. major*, leptomonad markers were expressed in these clusters at a higher level compared with the same markers in the procyclic forms (cluster 2). Unlike *L. major*, no clear early metacyclic population was apparent, with these markers being expressed in the late metacyclic (cluster 6) and in subpopulations of amastigote and procyclic clusters, mostly due to the presence of some proliferation makers in this set of genes. Finally, an unsurprising difference in the *L. mexicana* data compared to *L. major* was the ready detection of a discrete amastigote population (Fig. 4C, D).

The presence of *L. mexicana* late metacyclics and late leptomonads/haptomonads in the *in vitro* samples allowed us to ask if they arise from two distinct developmental trajectories, as was recently described for *L. major* in the sand fly^13^. To this end, we performed trajectory inference and pseudotime analysis to explore the development revealed by gene expression changes (Fig. 5). Strikingly, a branched trajectory was predicted, with a common developmental pathway from procyclics (cluster 2) through nectomonads/leptomonads (clusters 4 and 3_0) but splitting in the leptomonads (cluster 3_0) and then leading either to late leptomonads/haptomonads (cluster 3_1) or late metacyclics (cluster 6) (Fig. 5A). Differential expression analysis of genes across pseudotime was consistent with such forked development, with both inferred developmental lineages initially sharing changes in transcript levels but deviating around the inferred position of leptomonads and showing distinct patterns of expression thereafter (Fig. 5B-F). These data demonstrate that the branched developmental pathway of *L. major* in the sand fly^13^ is detectable in *L. mexicana* grown in culture, indicating it is likely a conserved feature of the *Leishmania* promastigote life cycle.

**Figure 5.**
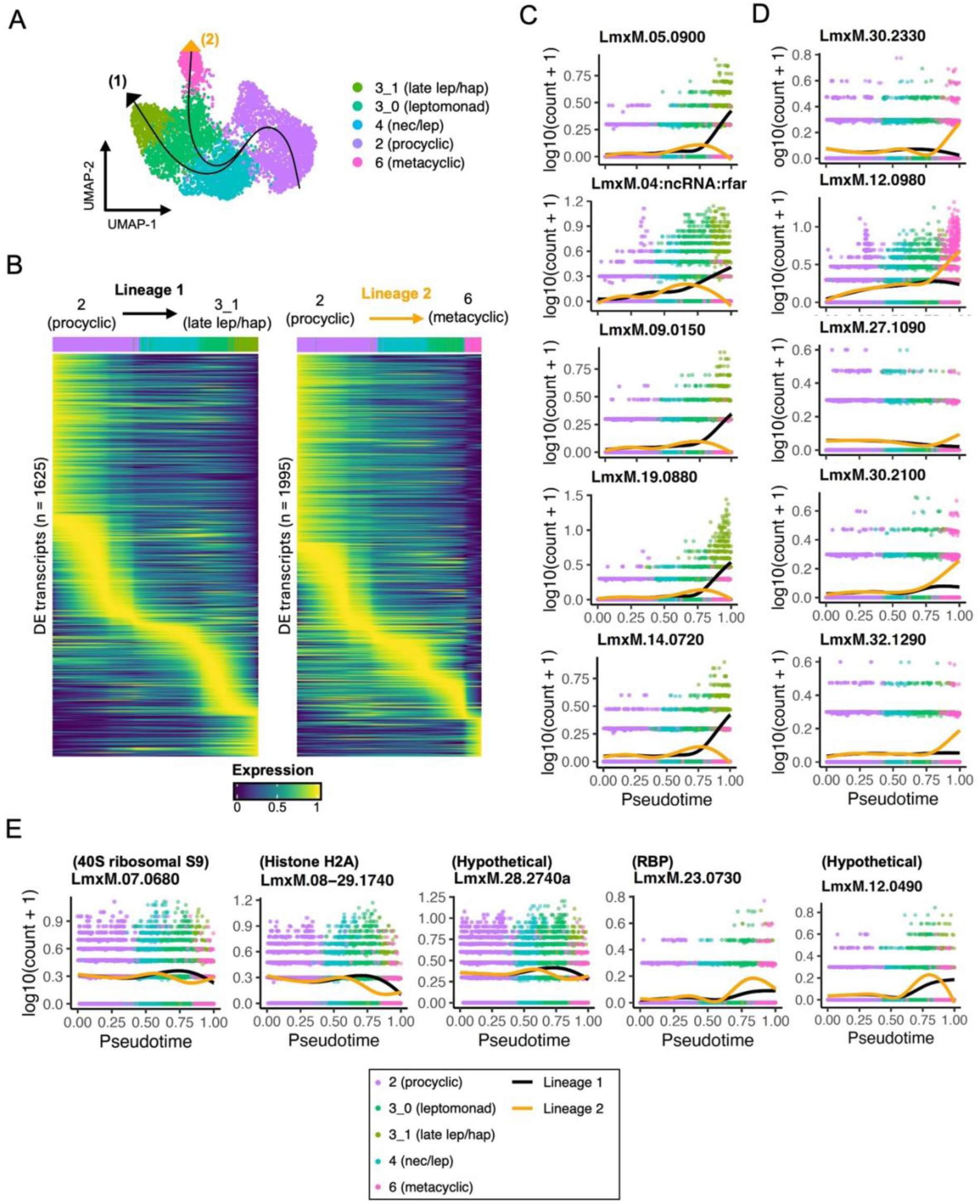
The developmental trajectory of *Leishmania mexicana* promastigotes in culture. **A)** UMAP coloured by cell type with branched predicted trajectory of development from procyclic to haptomonad (lineage1) and procyclic to metacyclic (lineage2). **B)** Heatmaps coloured by scaled expression for genes differentially expressed over lineage 1 (left) and lineage 2 (right). Scatter plots show the raw transcript counts of markers define the end points of lineage 1 **(C)** and lineage 2 **(D)**, as well differing over the point of divergence between the two trajectories **(E)**.

Of the genes with altered transcript levels over pseudotime (Fig. 5B), 1,219 were shared across the development, and 406 and 736 genes were uniquely differentially expressed in the lineages ending in late leptonomads/haptomonads (lineage 1) or late metacyclics (lineage 2), respectively. We first investigated the genes that differed between cells at the end of either trajectory. For the late leptomonad/haptomonad development the top five genes were LmxM.05.0900, a putative surface protein, a non-coding RNA encoding gene (LmxM.04:ncRNA:rfamscan:50220-50294), two ubiquitin-like proteins (LmxM.09.0150, LmxM.19.0880) and a hypothetical (LmxM.14.0720) (Fig. 5C). For the metacyclic lineage end point the top five were LmxM.30.2330, a putative 3,2-trans-enoyl-CoA isomerase, promastigote surface antigen PAS2 (LmxM.12.0980), a putative transporter (LmxM.32.1290)^40^ and two proteins of unknown function (LmxM.27.1090, LmxM.30.2100)) (Fig. 5D). These genes represent markers distinguishing late leptonomads/haptomonads from late metacyclics.

We next mined the data for putative regulators that might be driving the formation of the two lineages, by comparing transcript levels of genes during pseudotime 0.6-0.9, where the split in trajectory takes places. Fourteen of the top twenty differentially expressed genes were ribosomal proteins that are characterised by a “dip” in expression in lineage 2, ending in metacyclic development (an example is shown in Fig. 5E). The remaining were copies of histone H2A (Fig. 5E), amastin, tubulin and a putative heat shock protein (LmxM.28.2770). Focusing on genes with high expression in lineage 1, leading to late leptomonads/haptomonads, uncovered ribosomal proteins, histones and tubulin (Supplementary data 3). For lineage 2, however, the top ten genes included four genes encoding components for the paraflagellar rod, a copy of GP63 (LmxM.10.0460), a hypothetical protein (LmxM.12.0490) and an RNA binding protein (RBP, LmxM.23.0730) (Fig. 5E). This RBP has been found to be required for infection and proliferation in mouse footpads in *L. mexicana* and therefore is likely required for the development of infective forms. Unfortunately, despite our experimental set up successfully generating axenic *L. mexicana* amastigotes from stationary phase promastigotes, we could not infer a convincing trajectory, or trajectories, that led from the promastigotes to the amastigotes due to a lack of parasites between these two extremes.

### The conserved cell cycle regulated transcriptome of procyclic form trypanosomatids

Having confidently identified procyclic forms in our mixed *L. mexicana* population (Fig. 4A), we isolated these and applied our newly identified *L. major* cell cycle phase markers (syntenic orthologues) to this population (Fig. 6A). This analysis was able to label G1, S and G2/M cells that could then be arranged into a circular trajectory with Cyclum (Fig. 6B), as performed with *L. major* (above). Differential expression analysis identified 1,464 genes with dynamic transcript level changes over the cell cycle (Fig. 6C, Supplementary data 2). This afforded us the opportunity to identify robust cell cycle-regulated genes across the two *Leishmania* species investigated here, and in our previously defined *T. brucei* procyclic form cell cycle transcriptome ^16^. As these datasets were collected in independent experiments with varying numbers of cells and efficiency of transcript capture, not all genes can be meaningfully compared. Therefore, we limited our analysis to the expression patterns of genes that were detected in at least 10% of the *T. brucei* and *L. major* data sets, and 20% of the smaller *L. mexicana* procyclic dataset. Raw fold-change comparisons also did not allow for meaningful comparisons between datasets; to address this issue, we scored each gene using a variety of criteria. These included p values, fold change and the gene being present in the top-ranked cell cycle regulated gene set within and between each species (all comparison and scores are available in Supplementary data 4). Setting a threshold for genes with a minimum of 8/17 positive criteria in every species resulted in a list of 467 conserved orthologs scoring highly in all three species. This corresponds to 571 *L. mexicana* genes, 529

**Figure 6.**
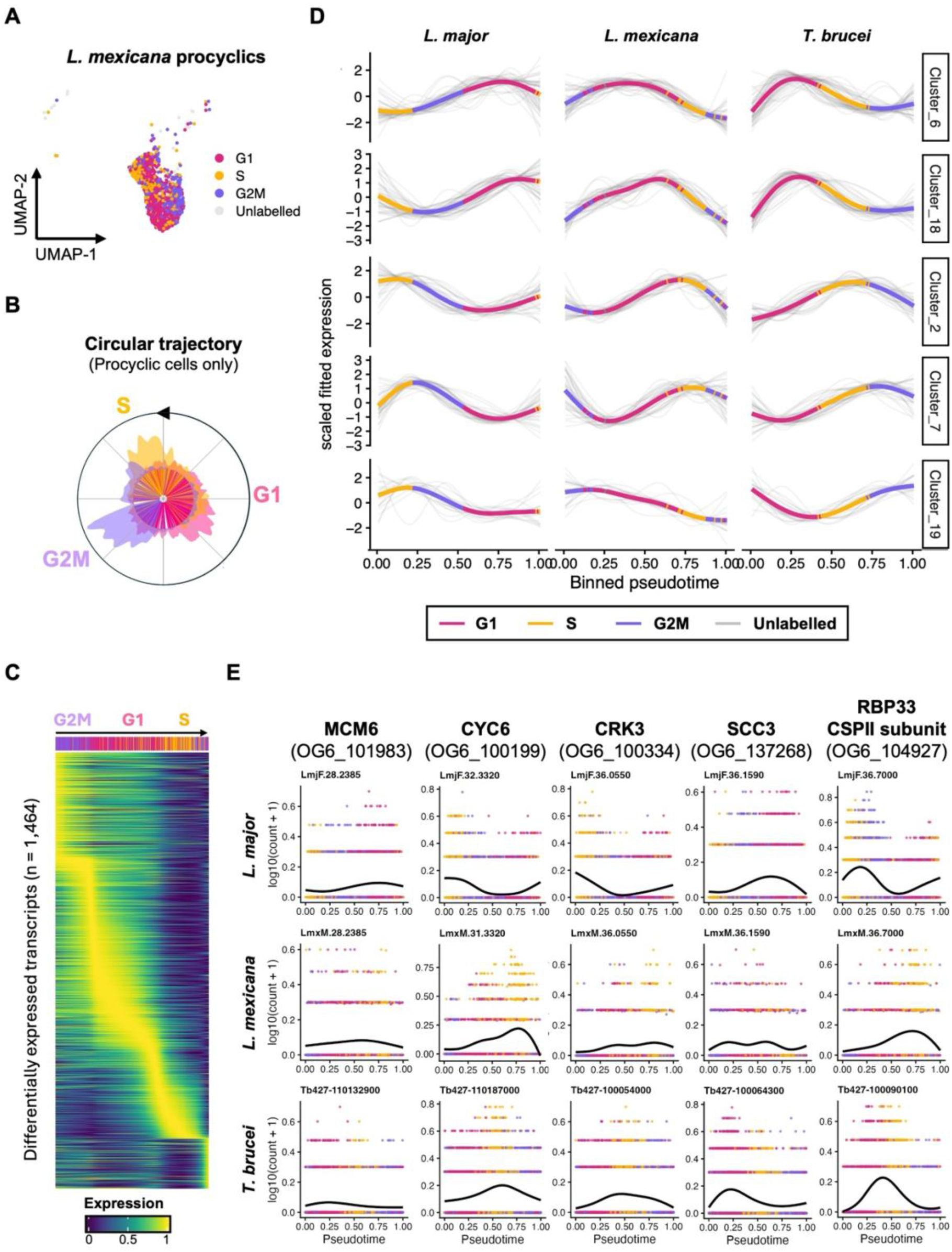
**Cell cycle regulated gene expression *Leishmania mexicana* and conservation with *L. major* and *T. brucei***. **A)** UMAP of Log phase *L. mexicana* procyclic promastigotes coloured by cell cycle phase. Phases were identified using markers identified in *L. major* procyclic forms. **B)** Cyclum generate circular trajectory, colours by phase. Centre is individual cells colours by phase, as in A), and outer ring shows histogram of the number of cells, split by phase. **C)** Heatmap showing scaled expression (z-score) of significantly differentially expression genes over the cell cycle. **D)** Example of co-expressed genes identified by hierarchical cluster, that shown conserved expression patterns across *L. major, L. mexicana* and *T. brucei*. **E)** Raw transcript counts of selected genes over pseudotime. Cells are coloured by phase and black lines shown average expression pattern.

*L. major* genes and 560 *T. brucei* genes due to some orthogroups containing multiple cell cycle regulated genes. We then used hierarchical clustering to group genes by expression pattern across the three species (Fig. 6D, Fig. S4, Supplementary data 4). Clusters were first defined using only 1:1 orthologs to allow direct comparison, and then remaining genes were assigned to the best matched expressed cluster.

Notably, not all clusters of orthologous genes peaked at the same precise point in the cell cycle of each species, but they did have conserved dynamics across species (Fig. S4, S5A). For instance, clusters 6 and 18 showed very similar conserved peak expression at the G1 phase in all species, providing 57-60 high confidence G1 marker genes for each species (Fig.6D). Clusters 2 and 7 peaked around S phase, and cluster 19 at G2/M, providing 63-64 and 22-25 marker genes, respectively, for these phases (Fig. 6D). Due to differing levels of detection between datasets, we did not aim to identify differentially expressed genes unique to each species, but it was notable that many of the clusters contained genes that, although cell cycle regulated in all three species, did not share conserved expression patterns (one example being a SET domain containing protein, Fig. S5B). Amongst known machineries and factors that act at specific stages of the cell cycle, the Origin Recognition Complex (ORC) and the Minichromosome Maintenance (MCM) helicase are widely conserved eukaryotic complexes that interact to bind and demarcate genomic loci as sites of DNA replication initiation^41^, termed origins. Although we now know most of these factors are generally conserved across kinetoplastids, these have been mostly studied in *T. brucei*^42–44^, while studies in *Leishmania*^45^ have reported mostly on one subunit of ORC and the dynamics of their expression across the cell cycle remains unknown in these parasites. Transcript levels of several of these factors were detected in our analysis (Fig. 6E, Fig S5A), including MCM5 (Fig. 6E), 6 and 7 (Fig. S5A) which were consistently expressed the highest during G1 or the G1 to S transition, suggestive of upregulation in their expression in preparation for S-phase. Most of the ORC subunits were not detected in all analysed datasets, with the exception of a putative ORC2 orthologue, which showed no cell cycle regulation (Fig. S5A). Notably, only two cyclins, CYC6 and CYC8, and two cdc-related proteins, CRK2 and CRK3, were detected and had conserved expression patterns in all three species (Fig. 6E, Fig. S5). CRK2 peaked in expression during late G1 (Fig. S5), in keeping with its essential role in S phase in *T. brucei,* where it phosphorylates MCM3 to promote its recruitment to the GINS complex at the origins of replication^46^. CRK3 transcripts peaked in S phase (Fig. 6E), again preceding the phase when this protein is known to function: CRK3 is required for the correct passage across the G2/M checkpoint in *L. mexicana*^47^ and *T. brucei* procyclic forms^48^. Similarly, CRK3’s binding partner, cyclin 6^49^, also peaked in S phase (Fig. 6E). A consistent G1 peak distribution was also seen for SCC1, SCC3, and SMCs 1, 2 and 4 (Fig. 6E, Fig. S5), which in other eukaryotes form the cohesin complex that plays roles in chromosome segregation and DNA repair^50^, including in *T. brucei*^51^, where the SCC1 protein is expressed from late G1-G2^52^.

Among several interesting orthogroups, Orthogroup OG6_104927 showed well-conserved expression (Fig. 6E) and, in *Crithidia fasciculata,* it binds two other subunits to form the cycling sequence binding protein 2 (CSBPII) complex that binds cycling transcripts ^53^. This provides one example of where the conserved ortholog list is enriched for cell cycle regulated markers, but where functional studies are frequently limited to one species. Therefore, the conserved expression patterns we describe suggest conserved functions across *Leishmania* and *Trypanosoma brucei*, and perhaps across wider trypanosomatids. Furthermore, the complete list of conserved cell cycle regulated genes includes several hypothetical proteins, hinting at undiscovered cell cycle biology perhaps unique to the trypanosomatids.

### Linking cell cycle and life cycle transitions in *L. mexicana*

Finally, we exploited the above list of highly conserved cell cycle-regulated genes to find the average expression of each across our complete *in vitro L. mexicana* datasets, encompassing all life cycle forms (Fig. 7A-C), akin to the analysis performed for *L. major* above. Comparing average expression of the cell cycle phase markers and life cycle markers (Fig. 7A-C) showed that the procyclic cluster 2 encompasses cells in G1, S and G2/M phases, with few (∼15%) unlabelled cells (Fig. 7B), consistent with these being cycling cells. Regarding the amastigote population, this analysis now offers an explanation for it being split into two clusters (0 and 5).

**Figure 7.**
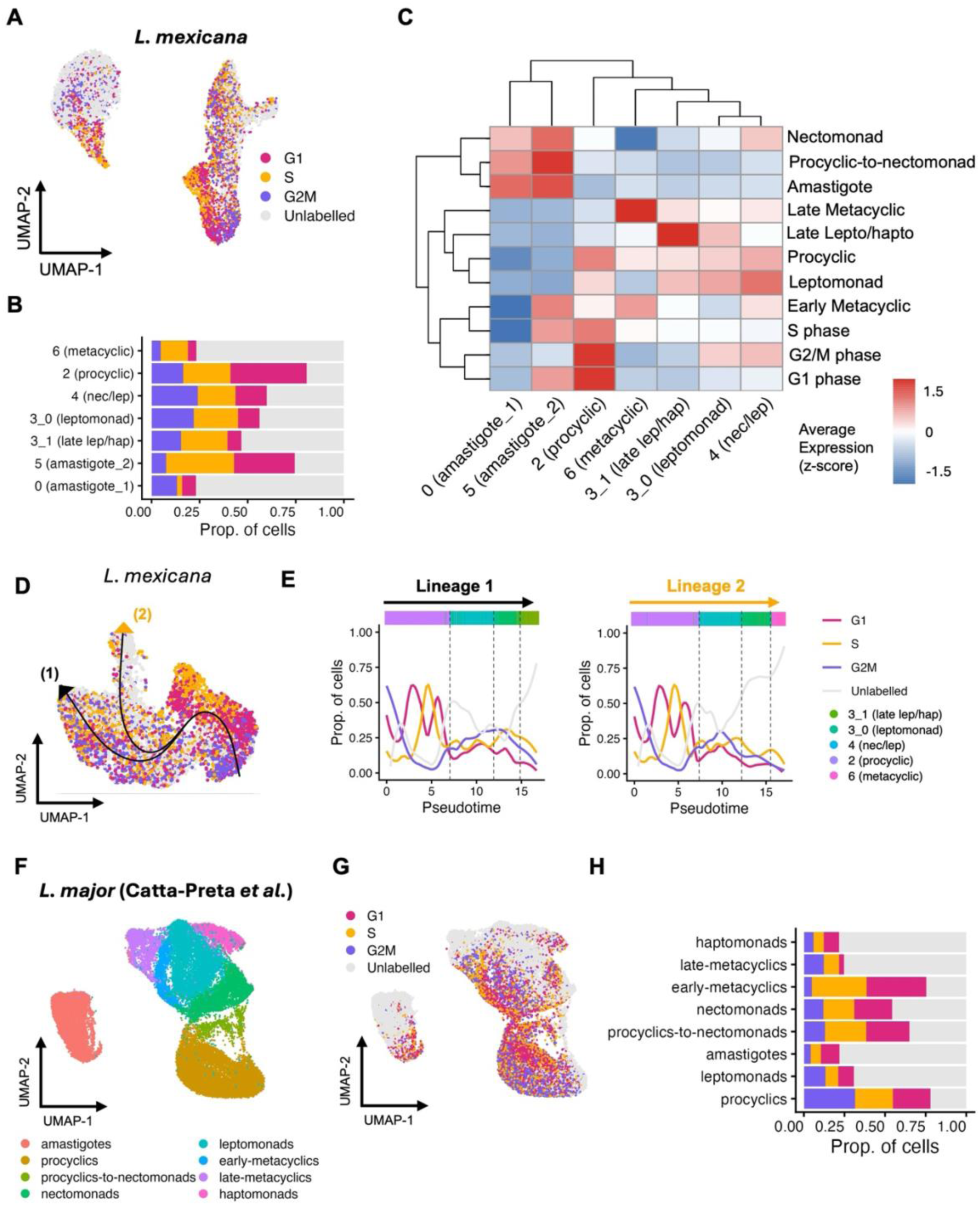
Identifying the cell cycle and life cycle transition links in *Leishmania* development *in vitro* and *in vivo*. **A)** UMAP of complete *L. mexicana* datasets coloured by cell cycle phase inferred using conserved marker set. **B)** Proportion of each clusters in each cell cycle phase or Unlabelled. Colours as in A). **C)** Scaled expression scores for both life cycle scores and cell cycle phase scores, for each cluster, with clustering to identify relationships. **D)** UMAP of promastigote cell types only, colours by cell cycle phase, with branched trajectory indicated. **E)** Proportions of cells in each phase across pseudotime for lineage 1 ending in haptonomands (left) and lineage 2 ending in metacyclics (right). F) UMAP of *L. major* data generated by Catta-Preta *et al.* for reference, clusters and descriptions are retained from authors original analysis. **G)** UMAP of in vivo *L. major* data colours by cell cycle phase. H) Proportion of cells in each cell cycle phase for clusters show in F.

Amastigote cluster 5 (amastigote_2) expressed similar numbers of G1 and S phase cells but G2/M phase expression was less evident (Fig.7A-C); this could be either due to low replication of this amastigote sample, or differences in the transcriptional signature of G2/M phases between the insect and mammalian infective forms of *L. mexicana*, as previously demonstrated in *T. brucei* ^16^. The other amastigote subcluster (Cluster 0, amastigote_1) was predominantly unlabelled (Fig.7A-C), perhaps due to early sampling after moving promastigotes into differentiation medium. Only approximately 25% of metacyclic cells (cluster 6) expressed cell cycle phase signatures, while the remaining cells were classed as unlabelled and are likely non-cycling (Fig.7A-C). Overall, this may suggest that cluster 6 is a mix of early replicating, and late arrested metacyclics, as expression of both early and late metacyclic marker sets was detected (Fig. 7C). However, the low number of cells in this cluster prevented separation of these early and late forms. Of the remaining cells expressing nectomonad, leptomonad and haptomonad signatures (clusters 3_0, 3_1 and 4, Fig. 7A), approximately 50-60% of each population expressed cell cycle phase markers, with the rest being unlabelled (Fig. 7B, C). Plotting the change in cell cycle phase over the branched promastigote developmental trajectory (Fig. 7D, E) indicated nectomonad/leptomonad cluster 4 cells emerge from G1 procyclics, and that both late leptomonad/haptomonad and metacyclic end populations of each lineage are enriched with non-cycling cells. However, the cycling and non-cycling cell proportions present across the nectomonad and leptomonad developmental stages showed variation between the two development linages: in the leptomonad cluster 3_0, development to metacyclics was linked to a higher proportion of unlabelled cells (∼70%) compared to those in the haptomonad developmental pathway (∼30%) (Fig. 7E). These data reveal that cell cycle exit occurs in both branches of development from leptomonads but suggest that the dynamics of these developmental changes leading to metacyclics and haptomonads are distinct.

To ask if these dynamics of *Leishmania* cell cycle and life cycle progression *in vitro* reflect that of promastigote development in the sand fly, we took the complete dataset generated by Catta-Preta *et al.* ^13^(Fig. 7F) and similarly applied our optimised cell cycle phase labelling method (Fig. 7G). Labelling of this dataset using only G2/M phase markers that are conserved across the three trypanosomatids reported in this study led to underestimation of this population of cells (Fig. S6), which is consistent with species-specific differences at this cell cycle phase. To circumvent this, *L. major* G2/M markers derived from our *in vitro* dataset reported above were used (Fig. 7G). This analysis showed that the pattern of cell cycle change within promastigote development *in vivo* (Fig. 7G, H) closely matches the patterns of *L. mexicana* development *in vitro* (Fig. 7B, C). *L major* late metacyclics and haptomonads were largely unlabelled (and so non-cycling), whereas procyclics and early metacyclics were cycling (Fig. 7G, H). In each case, the proportions of unlabelled or cell cycle phase labelled cells mirrored that seen for *L. mexicana in vitro* (Fig. 7B, C). Intriguingly, G2/M phase cells were more abundant in the *in vivo L. major* late metacyclics, with early metacyclics being enriched for G1 and S phase cells (Fig. 7H), an observation we also made in our *L. major in vitro* data (Fig.3D), indicating that the early to late metacyclic transition occurs in G2/M prior to the final cell division. Again, nectomonads and leptomonads did not separate cleanly as non-cycling and proliferative populations, respectively, as would be predicted by early morphology classification studies (Fig. 7G, H)^4,7^. Instead, this analysis suggests a mix of arrested and proliferating cells in both cell types, in keeping with Catta-Preta and colleagues’ earlier observations of dispersed cell cycle phases in these forms^13^, and our observations of *L. mexicana* development *in vitro*.

Together, these analyses demonstrate that promastigote life cycle developmental transitions are each linked to varied cell cycle dynamics: procyclic to nectomonad/leptomonad development appears to occur during G1, whereas subsequent progression to arrested haptomonads is not clearly linked to a defined intra-G1 developmental switch; in contrast, metacyclic development involves an intermediate proliferative form, with G2/M being the terminal phase before cell cycle arrest.

## Discussion

In this work, we have applied scRNA-seq to address two aspects of *Leishmania* biology in *in vitro* culture: the presence or absence, and gene expression profile, of known developmental forms seen *in vivo,* and changes in gene expression across the cell cycle of replicating life cycle stages. Such analyses are dependent on the depth of information that is provided by scRNA-seq, a methodology that does not rely on isolation of life or cell cycle forms^12^, and expands our capacity to interrogate and understand the complex development of *Leishmania* cells across the life cycle. A key finding of our work is that *L. mexicana* grown *in vitro* represent a strong model for the developmental pathways of *Leishmania* seen in the sand fly: not only could we see comparable gene expression in distinct populations representing most of the life cycle forms that have been described *in vivo*, we were also able to show that the branched pathway of *L. major* development leading to either metacyclics or haptomonads^13^ was recapitulated in culture (Fig. 5). Perhaps surprisingly, we did not find evidence for comparable development in *L. major* in culture (Fig. 3), as we could not detect clear leptomonads or haptomonads. It is possible that the lack of these forms merely reflects the fact that we sampled *L. major* only during exponential growth or in early stationary phase, thus failing to capture these populations, perhaps because they accumulate at prolonged higher densities in culture (which may explain why they were detected in our *L. mexicana* experiments). Alternatively, replicative leptomonads may simply not survive or arise in our culture conditions, in contrast to the sand fly environment. These data may reveal why *L. mexicana* has been the better model for promastigote to amastigote development *in vitro*^39^. The ability of scRNA-seq to identify markers of late metacyclic, leptomonads and haptomonad forms provides a means to test conditions that might allow the branched terminal development to occur in *L. major* growing in culture, expanding the capacity for *in vitro* dissection of life cycle development beyond *L. mexicana.* Despite the absence of detectable leptomonads and haptomonads *in vitro*, scRNA-seq shows that stationary phase *L. major* promastigotes are transcriptionally highly distinct from exponentially growing promastigotes and do not exclusively comprise non-replicating cells (Fig. 1, 2). Within these stationary phase cell types, we could readily detect both *L. major* early metacyclics and late metacyclics, as seen *in vivo* ^13^. The presence of metacyclics suggests that the development of these forms, at least in culture, is not dependent on the other branch (which leads to haptomonads). However, the absence of detectable leptomonads is unexpected, as scRNA-seq of *L. major in vivo*^13^ and *L. mexicana in vitro* (this study) indicates these are the progenitors of both haptomonad and metacyclic non-replicating life cycle forms. Although trajectory inference was not employed in the case of *L. major* promastigote development (due the negligible presence of cells between the two large populations limiting biological interpretation), the early metacyclic cluster positions close to late metacyclics, suggesting these late forms do develop from the early metacyclics, as observed *in vivo*. The early stages of development that give rise to the other STA subpopulations are less clear-cut. Early metacyclics may arise by a complex route from procyclics or nectomonads without the need for cycling leptomonads as intermediates, at least *in vitro*. If so, this may indicate flexibility in life cycle development, as has been debated for *T. brucei*^54–56^. In fact, even the Log sample of

*L. major* contains low numbers of non-proliferative clusters 1 and 3, and early replicating metacyclics, suggesting only late metacyclics are exclusive to higher density stationary phase cultures. These data may indicate that the pathway for strict ‘normal’ development is lacking in the *L. major* cells we have used relative to Catta-Preta and colleagues, perhaps due to prolonged *in vitro* culture and akin to the ‘monomorphism’ seen in some *T. brucei* BSF cells^57^.

Based on the expanding scRNA-seq datasets available for trypanosomatids, a consensus may be emerging in the formation of trypanosomatid life cycle forms that are truly non-cycling and act as transmission forms: in each case, the development of these cell types is preceded by, as yet poorly-defined, transitional cell types that share transcriptional signatures with their successors. Here, as was seen *in vivo*^13^, *L. major* and *L. mexicana* non-replicating late metacyclics arise from replicating early metacyclics (Fig. 3D, 7C). This developmental programme bears comparison with the formation of mature *T. brucei* metacyclics in the salivary gland of the tsetse fly, which are preceded by pre-metacyclics^58,59^, with the two forms differing in expressing a single or multiple VSGs, respectively^60^. Indeed, a further similarity is seen in *T. brucei* development in the mammalian bloodstream form, where cell cycle-arrested, morphologically stumpy form cells that act in transmission to the tsetse arise via the formation of cells with a stumpy-like transcriptome but that are not yet committed to cell cycle exit or have changed in morphology^17,61^. Similarly, recent scRNA-seq analysis of the *T. cruzi* life cycle has indicated the presence of sub-populations of cell types within life cycle forms found in both the vector and mammalian hosts, consistent with staged differentiation, but to date have not been linked to cell cycle dynamics^62,63^. Whether these similarities extend to conserved developmental control factors^17^ or modes of cell cycle commitment and exit^61,64^ requires further work.

At least two of the four *L. major* stationary populations we have described here are considered non-replicating life cycle forms (nectomonads and late metacyclics), whereas one (early metacyclic) appears to have G1 and S-phase characteristics and, as discussed above, is likely a transitional form. However, we have not been able to assign one other population (cluster 1) to a known life cycle form (Fig.3D). One possibility is that cluster 1 is leptomonad-like, as it expresses some leptomonad markers, but is not recognisable as such due to an inability to be established or sustained in culture, perhaps due to mutation, explaining the lack of cell cycle marker expression and haptomonads. Intriguingly, one of the top specific markers for cluster 1 is GCS1/HAP2 (Lmj.35.0460) (Fig. S1B, arrow), which is highly expressed in the novel cell populations that arise after g-irradiation treatment of *L. tropica*, and has been shown to be needed for *Leishmania* mating^18,37^. Akin to treatments with other DNA damaging/stressing agents, g-irradiation treatment of *L. tropica* and *T. cruzi* leads to an increase in mating between vector life cycle forms of these parasites in culture ^18,65^. Unlike *L. tropica*, no scRNA-seq has been done in *T. cruzi* post g-irradiation treatment, but it is possible that in both parasites DNA stress enhances a natural process of cell cycle exit, which is distinct from the formation of haptomonads or metacyclics in *Leishmania*, and enriches for so-far poorly understood trypanosomatid populations that act in mating. Whether this is also seen in *T. brucei* is unclear, since *in vitro* mating of both *L. tropica* and *T. cruzi* results in whole genome tetraploid cells, which appear distinct from *T. brucei* meiosis in the tsetse fly that results in the formation of haploid gametes^66^, despite these also expressing HAP2^59^.

Akin to our previous work in *T. brucei*^16,17^, we have been able to detect a large set of genes whose transcript abundance varies across the cell cycle in both *in vitro Leishmania* scRNA-seq datasets (Fig. 2, 6). Comparing the cell cycle-regulated transcriptomes of the two genera revealed a number of conserved functions, in particular in G1 and S phases, as well as many unexplored hypothetical genes (Supplementary data 3). A limitation of 10X-based scRNA-seq is partial recovery of the transcriptome in single cells, meaning that the identification of species or genera-specific cell cycle expression can be problematic and is inevitably incomplete.

Notably, fewer conserved G2/M phase genes were common to the three datasets, perhaps suggesting that this stage of the cell cycle provides greater opportunity for flexibility across the three examined species. In this regard, it is known that distinct post-G2 cell cycle checkpoints are present in bloodstream and procyclic forms of *T. brucei*^48,67,68^, and that bloodstream form cells can continue to divide in the presence of unrepaired DNA breaks^69^. *Leishmania* is known to display mosaic aneuploidy^70^, which may relate to progress through mitosis. Indeed, such innovation may not be limited to trypanosomatids, since schizogony during the life cycle of the malaria parasite, *Plasmodium*, likely requires modulation of a checkpoint between mitosis and cytokinesis^71,72^. As discussed above, cell cycle exit is key to parasite life cycle differentiation and, in this regard, scRNA-seq shows that late metacyclics in both *L. major* and *L. mexicana* appear to arrest at the G2/M stage of the cell cycle, unlike *L. mexicana* haptomonads, suggesting variation in checkpoint use during exit. In fact, life cycle transitions may display even greater variation in cell cycle controls, since nectomonads, which are thought to be non-cycling^4^, did not show clear evidence for cell cycle arrest in either *in vitro* (this study) or *in vivo* scRNA-seq datasets^13^.

Perhaps these life cycle forms, unlike metacyclics, are only partly arrested, for reasons that remain unclear. Understanding how ‘true’ arrested forms restart the cell cycle, such as when metacyclics enter the mammal, may shed light on the biology of growth resumption, but such transitions have not yet been captured by scRNA-seq analyses.

## Methods

### Parasite culture

*Leishmania major* Friedlin and *L. mexicana* M379 promastigotes were cultured in haemoflagellate minimal essential medium (HOMEM) supplemented with 10% heat inactivated foetal bovine serum (FBS), at 25 °C in non-vented flasks. For differentiation to amastigote forms, late log/stationary phase *L. mexicana* promastigote cultures were collected by centrifugation and then resuspended at ∼1 x 10^6^ cells/ml in Scheider’s Drosophila Medium at pH 5.5, supplemental with 20% FBS and 15 mg/ml of haemin, and maintained at 32 °C with 5% CO_2_ in vented flasks.

### Flow cytometry analysis

For flow cytometry analysis of DNA content by propidium iodide (PI) staining, 6 x 10^6^ *L. major* promastigote cells were collected into 15 ml tubes and processed using a standard cell cycle profile analysis protocol with few modifications as described before^73^. Briefly, the cells were centrifuged at 1000 g for 5 min, washed in 1 ml of filtered 1x PBS (Sigma-Aldrich) supplemented with 5 mM of EDTA (Gibco) and 1% Fetal Bovine Serum (Gibco) (PBS-EDTA-FBS), centrifuged at 1000 g for 5 min, and re-suspended in 900 ml of PBS-EDTA-FBS by careful pipetting. Next, 2.1 ml of 100% Methanol (stored at -20°C) were added dropwise (final concentration of 70% methanol) while gently vortexing/swirling the tube, which was then stored at 4°C overnight protected from light. The cells (1 ml; ∼2 × 10^6^ cells) were then centrifuged at 1000 g for 5 min, washed once in 1 ml of PBS-EDTA-FBS, and gently re-suspended in 300 µl of PBS-EDTA-FBS supplemented with 10 µg/ml of Propidium Iodide and 20 µg/ml of RNase, after which they were incubated for 45 min at 37°C protected from light. Next, the cells were transferred into 6 ml tube (BD Falcon™ tube,BD Biosciences) through a mini cell strainer (pluriStrainer) with a mesh size of 40 µm and run on the University of Glasgow MVLS Shared Research Facilities BD FACSCelesta Multicolour Flow Cytometer (BD Biosciences). Data was acquired from the FSC, SSC and PE channels using the BD FACSDiva software (BD Biosciences). Data was then analysed using FlowJo (v10).

### Single cell RNA-seq sample preparation and sequencing

For both *L. major* and *L. mexicana*, samples were prepared and processed for 10X Genomics Chromium scRNA-seq as described previously in Briggs *et al*., 2023 and Briggs *et al*., 2021, respectively. In the case of *L. major*, 8,000 cells of each of the Log and stationary (STA) cultures (Fig. 1A) were mixed with 16,000 *T. brucei* bloodstream and procyclic form cells into two samples, 1 and 2 (the *T. brucei* samples and analysis have been published^16^). For *L. mexicana*, five staggered experimental cultures (three promastigote and two axenic amastigotes) were prepared in order to collect the five samples at the same time point. For replicate 1 (rep 1, Fig. S3D), 3,000 cells from each culture (total of 15,000 cells) were multiplexed with *T. brucei* bloodstream form cells into one sample (sample 1) (the *T. brucei* samples and analysis have been published^17^); for replicate 2 (rep 2, Fig. S3D), the five cultures were split into two samples, one containing promastigote cells (sample 2 Pro, Fig. S3E) and another containing amastigote cells (sample 2 Ama, Fig. S3E). Sample 2 Pro was multiplexed with *T. brucei* cells, while sample 2 Ama was multiplexed with *T. congolense* bloodstream forms.

### Read mapping, quality control and preprocessing of scRNA-seq data

Read mapping and transcript counting was performed with Cell Ranger version 7.0.0^74^ using “threeprime” chemistry and setting r1 length to 26. Mapping to both *Leishmania* and *Trypanosoma* references simultaneously - “chimeric reference genomes” generated using Cell Ranger’s mkref function - allowed for downstream demultiplexing of each species and removal of cross contaminating transcripts. For *L. major*, the *L. major* Friedlin nuclear reference genome^75^ (TriTrypDB v50) and *Leishmania braziliensis* maxicircle kDNA sequence (GenBank: LR697134)^76^ were combined with *T. brucei* Lister 427 2018 reference genome^77^. For *L. mexicana*, the *L. mexicana* MHOM/GT/2001/U1103 nuclear reference genome (TriTrypDB v55)^78^ and the *Leishmania braziliensis* maxicircle kDNA sequence (GenBank: LR697134)^76^ were combined with the *T. brucei* TREU927 nuclear reference genome (TriTrypDB v50)^79^ and kDNA maxi circle reference genome (GenBank: M94286.1). For the *L. mexicana* samples multiplexed with *T. congolense,* the *L. mexicana* genome (above) was combined with the *T. congolense* IL3000 2019 reference nuclear genome^80^. To allow mapping to the 3’ end of genes in the absence of high confidence annotations, in all cases the 3’ UTR annotations were extended to 2,500 bp or until the next genome feature annotation to avoid overlaps, as performed previously^17^. In case of *L. mexicana* samples, DecontX^81^ was used as an additional quality control to remove any contaminating free transcripts with default parameters. Cross-species mutliplets were manually removed by sample visualisation with tSNE and residual cross contaminating transcripts were then removed from single species transcriptomes. Finally standard quality control (QC) was performed to remove transcriptomes with below or above average total transcripts per cell, total features (genes) captured per cell, above average percentage of kDNA mitochondrial transcripts and rRNA transcripts. Post-QC, 4,169 and 4,009 transcriptomes were kept for *L. major* samples 1 (Log) and 2 (STA), while 6,192, 8,308 and 8,250 transcriptomes were retained for *L. mexicana* sample 1 (combined promastigote and axenic amastigote samples), sample 2 promastigote-only and sample 2 amastigote-only conditions. All code and cellranger count outputs are available at Zenodo 10.5281/zenodo.23016438.

### scRNA-seq data merging, integration, dimensional reduction and clustering

In all cases, counts were log2 normalised with Scran and scater^82^ packages before replicate two samples (promasitgote-only and amastigote-only) were simply merged with Seurat version 5.1.0^83^ as sample preparation and sequencing was performed in a single batch. Replicates 1 (rep 1) and 2 (rep 2) were then integrated using the CCA integration method implemented in Seurat. UMAP dimensional reduction was then performed as described in Briggs *et al*., 2021 and 2023: for *L. major*, 9 dimensions and 0.3 resolution was used; for *L. mexicana*, 20 dimensions were used, with clusters being identified with a resolution of 0.25, while subclusters for the leptomonad/haptomonad population were identified using a resolution of 0.2. *De novo* marker genes for clusters were identified with FindAllMarkers function using MAST^84^ for differential expression between clusters.

### Trajectory inference and differential expression analyses

For trajectory inference of *L. mexicana* promastigote development, Slingshot^85^ was applied with default settings without defining starting or ending clusters. To capture the cell cycle, Cyclum periodic trajectory inference ^34^ was performed as previous described^17^. To enable differential expression analysis across pseudotime with PseudotimeDE^35^, trajectories were inferred for 100 subsets of cells (80% of the population) in every case. PseudotimeDE was only used to test expression of genes detected in at least 10% of the target cell types, to avoid biasing analysis with genes with low detection across the population. Tests were applied to log2 normalised counts data using a gaussian model.

### Life cycle and cell cycle marker investigation

For life cycle marker investigation in *L. major* all cell type markers were taken directly from *in vivo* study by Catta-Preta *et al.* (2024)^13^; however; genes encoding ribosomal proteins were removed with the exception of the “procyclic to nectonomad” for which all but one marker was a ribosomal protein. For the analysis of the *L. mexicana* datasets, syntenic orthologs of these identified using OrthoMCL^86^ via TriTrypDB^87^.

The AddModuleScore function from Seurat was used with each list to find each marker lists average expression score per cell, defined as the average expression of the maker list minus the expression of a control feature set of genes. For cell cycle phase markers, two lists were used: first (Fig. 2A), marker genes from *T. brucei* were obtained from Briggs *et al*., 2023^16^ and a list of syntenic orthologs (excluding multi-copy genes) in *L. major* was created; second (Fig. 2E), cell cycle-regulated genes obtained from the analysis of the *L. major* sample (Fig. 2C ^13^) were further filtered (see results) to create a *Leishmania*-specific list. The MetaFeature function was used where the relative contribution of each marker list is calculated per cell. For phase labelling, the ratio of each score per cell compared to the average score across the sample was first calculated. The distributions of these ratios were used to guide the selection of a threshold of 1.05-fold change to assign a cell cycle phase per cell. f for a given cell the fold change was below 1.05 for every phase marker set (G1, S and G2/M) the cell was assigned as “Unlabelled”.

## Data availability statement

Raw sequencing data is available via ENA Project PRJEB127436. Processes data, objects and associated code are available via Zenodo (10.5281/zenodo.23016438). Results are available as supplementary data files.

## Acknowledgments

We thank J. Galbraith (Glasgow Polyomics, University of Glasgow) for library preparation and sequencing services, and all members of the McCulloch lab for discussions. This work was supported by the Wellcome Trust (224501/Z/21/Z to RM, 218648/Z/19/Z to EMB), the MRC (MR/Z504786/1, awarded to EMB), a Wellcome Trust Institutional Strategic Support Fund award held at the University of Glasgow (310182 to CAM), and the BBSRC (BB/N016165/1, BB/R017166/1 to RM, and BB/W001101/1 to RM and CAM). The Wellcome Centre for Integrative Parasitology was supported by core funding from the Wellcome Trust [104111].

## Author contributions statement

Conceived the study: CAM, EB, RB, TO, ML, RM. Designed and conducted the research: CAM, FW, GS, MY, EB. Analysed and interpreted the data: CAM, RL, FW, KC, RB, TO, ML, RM, EB. Wrote the initial draft: CAM, RM, EB. Edited and approved paper: CAM, RL, FW, KC, RB, TO, ML, RM, EB. Funding: EB, CAM, RB, RM, ML.

---

**Figure S1.**
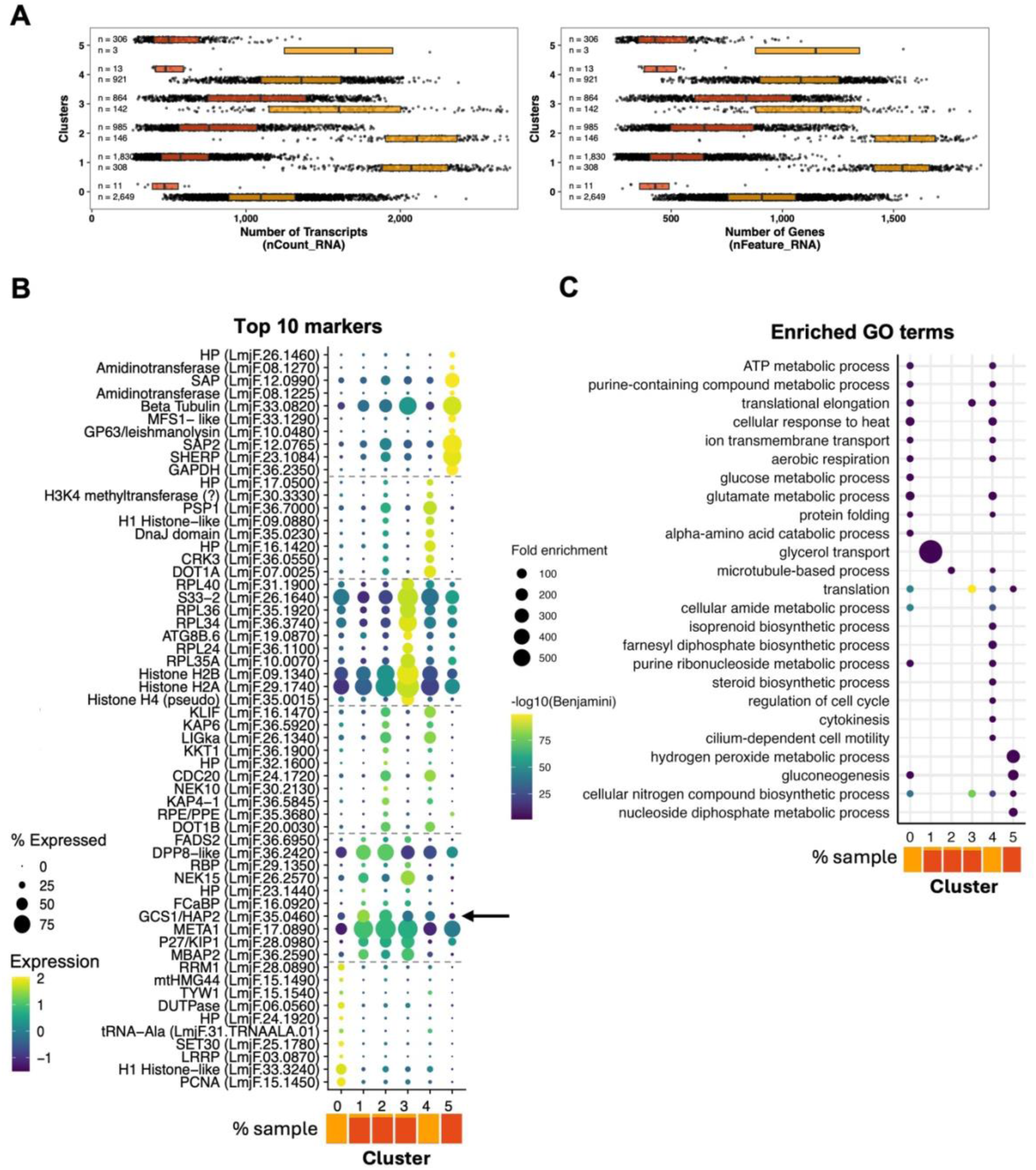
**A)** Number of total transcripts (left) and genes (right) detected per cell, separated by clusters. Box plot shows mean and interquartile range. **B)** Dotplot showing the top 10 markers for each cluster. Stacked bar chart below x-axis shows the proportion of each cluster that originated from each sample. **C)** Biological process GO terms enriched in the complete set of marker genes for each cluster.

**Figure S2.**
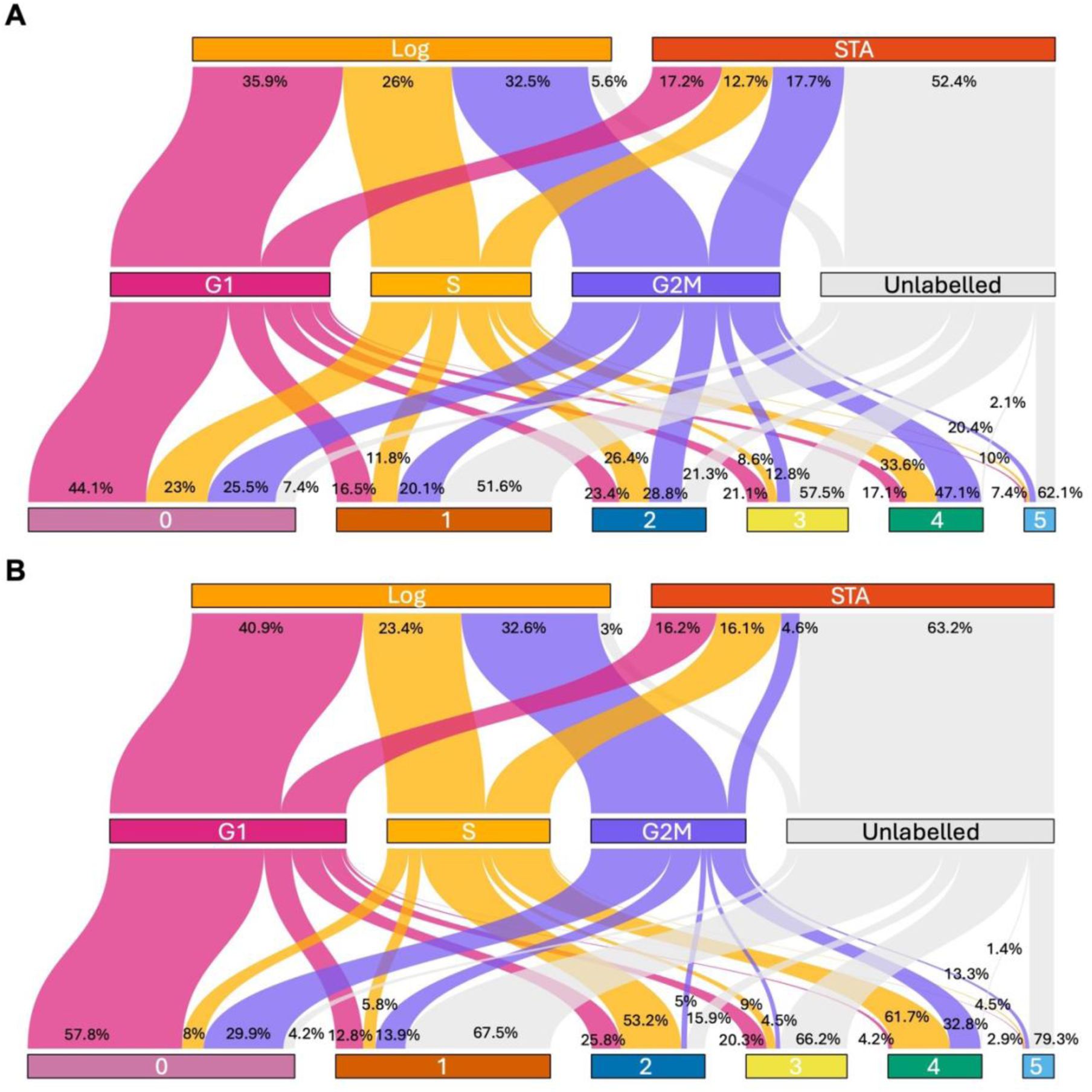
**A)** Alluvial plot showing the proportion of each sample (top) and cluster (bottom) labelled as each cell cycle phase (middle), defined using the *T. brucei* cell cycle markers. Relates to Fig. 2A. **B)** Zoomed plot from Fig. 2F, with the percentage of cells per sample/cluster labelled with the different cell cycle stages.

**Figure S3.**
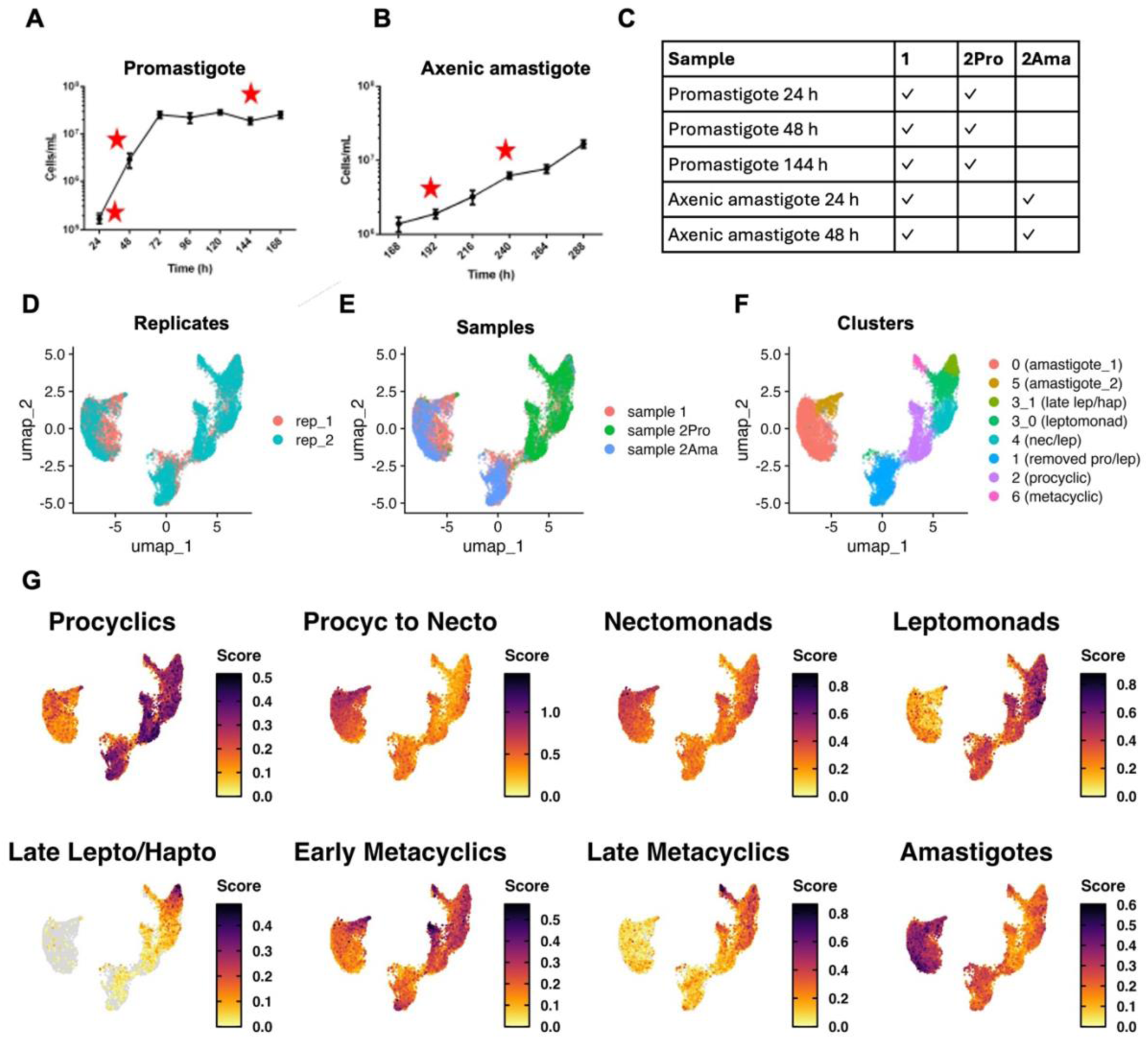
Growth curves of *L. mexicana* promastigote culture (**A**) and axenic amastigote culture (**B**). Red stars indicate time points sampled for scRNA-seq. **C)** Table indicating which timepoints were included in each sample. UMAPs of all samples coloured by biological replicate (**D**), samples (**E**) and clusters before the removal of cluster 1 (**F**). **G)** UMAPs all *L. mexicana* samples coloured by average expression scores of complete homologous marker list identified previously, for each life cycle form, before the removal of cluster 1.

**Figure S4.**
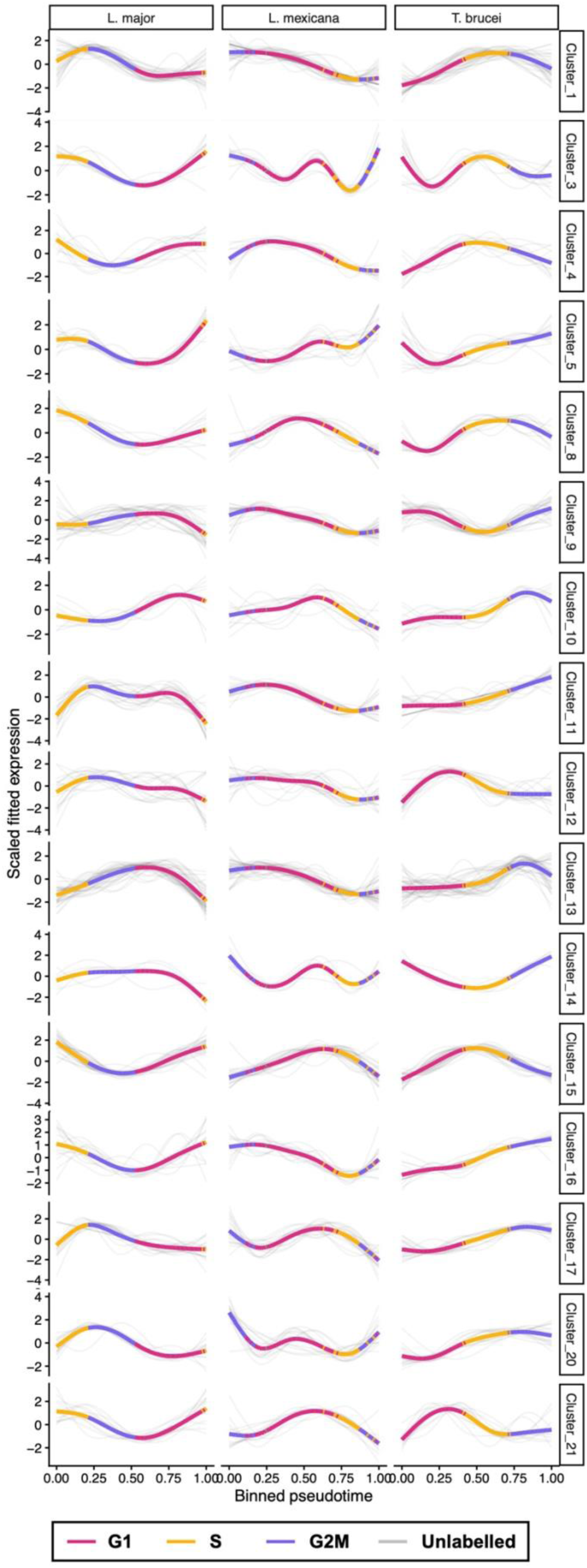
Plots of fitted expression all co-expressed genes identified by hierarchical cluster, that shown varied expression patterns across *L. major, L. mexicana* and *T. brucei*.

**Figure S5.**
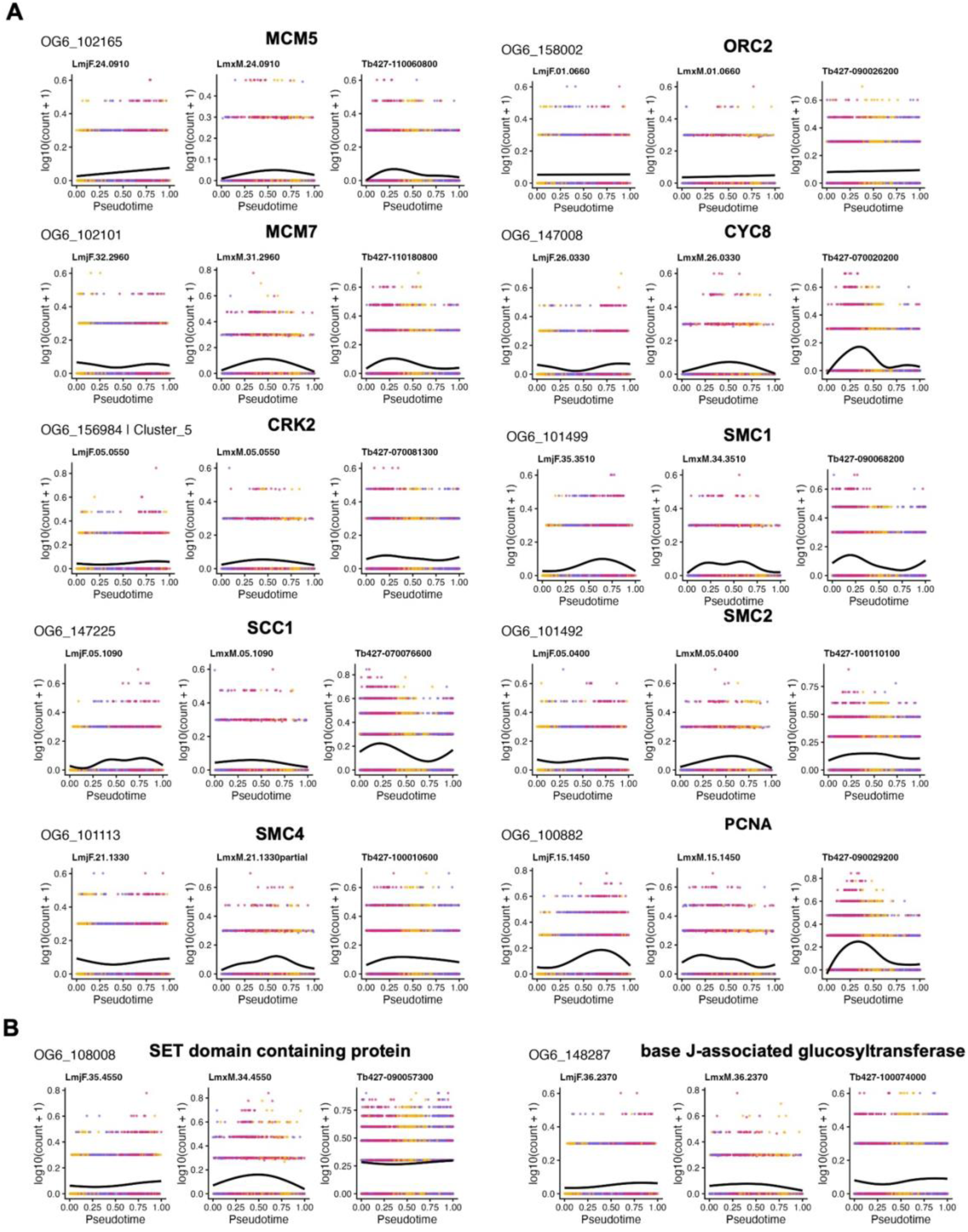
More examples to support Fig. 6E. Raw transcript counts of selected genes over pseudotime. Cells are coloured by phase and black lines shown average expression pattern.

**Supplementary data 1 –** De novo cluster marker discovery and associated GO terms for *L. major* promastigotes.

**Supplementary data 2 –** Differentially expressed transcripts over the *L. major* and *L. mexicana* procyclic cell cycles

**Supplementary data 3 –** Differentially expressed transcripts during *L. mexicana*promastigote *in vitro* development

**Supplementary data 4 –** *Leishmania sp.* and *T. brucei* cell cycle regulated transcriptome comparisons

